# Learning the limits of protein structure models with NMR chemical shifts

**DOI:** 10.64898/2026.11.12.766294

**Authors:** Cizhang Zhao, Gina El Nesr, Yilin Chen, Hannah K. Wayment-Steele

## Abstract

Protein foundation models are intended to implicitly learn the chemical rules underlying protein function. However, systematically evaluating what biochemistry these models have learned remains difficult, as most downstream tasks lack experimental data fine-grained enough to expose their failure modes. NMR chemical shifts fill this gap: as direct per-atom reporters of local chemical environment, they offer a sensitive probe of structure-based representations. In this work, we present a systematically curated chemical shift dataset designed to minimize experimental confounders. We use it to benchmark diverse protein foundation models in their ability to predict backbone ^15^N chemical shifts, atoms that are known to be sensitive to many effects. The largest error across all models is in core residues with evidence of fast-timescale dynamics, which we investigate mechanistically using molecular dynamics simulations. Given the biological relevance of structured dynamic regions, we trained a new model based on the best of these embeddings, SLAE, to predict its own error. “SLAE-ERR” achieves a mean absolute error of 0.76 ppm for predicting error. Together, we hope that the datasets, evaluation tasks, and models presented here are sources of guidance for protein design model improvement.

## Introduction

Capturing local chemical environments in protein structure is a foundational requirement for reliable performance across deep-learning-based protein modeling, such as protein structure prediction^1,2^, protein design^3^, protein property prediction^4,5^, and general structure encoding^6^. While substantial progress has been made in machine learning to improve the field’s predictive understanding of proteins^2,6^, comparatively little work has systematically examined what these models have learned about local chemical environments in proteins, in part because most downstream tasks lack the appropriate ground truth experimental data for benchmarking.

Nuclear magnetic resonance spectroscopy (NMR) enables solution measurements of protein structure^7^, interactions^8^, and dynamics^7,9,10^ under ambient conditions. NMR has been critical to understanding how enzymes perform catalysis^11^, drugs exert inhibition^12^, and mutations alter structure and function^13^. Its most common observable, the chemical shift, describes the local magnetic shielding of atoms, and therefore reflects the local chemical environment and subsequent perturbations resulting from protein function or dynamic processes^14,15^ (Fig. 1a, b). Therefore, chemical shift prediction is a direct, per-atom prediction task that evaluates how well structure-based models capture local chemical environments^5,6^. Accurate chemical shift prediction is useful in its own right for downstream tasks such as NMR assignments^16,17^, structure determination^18^, and inference-time sampling of protein generative models^19^.

**Fig. 1.**
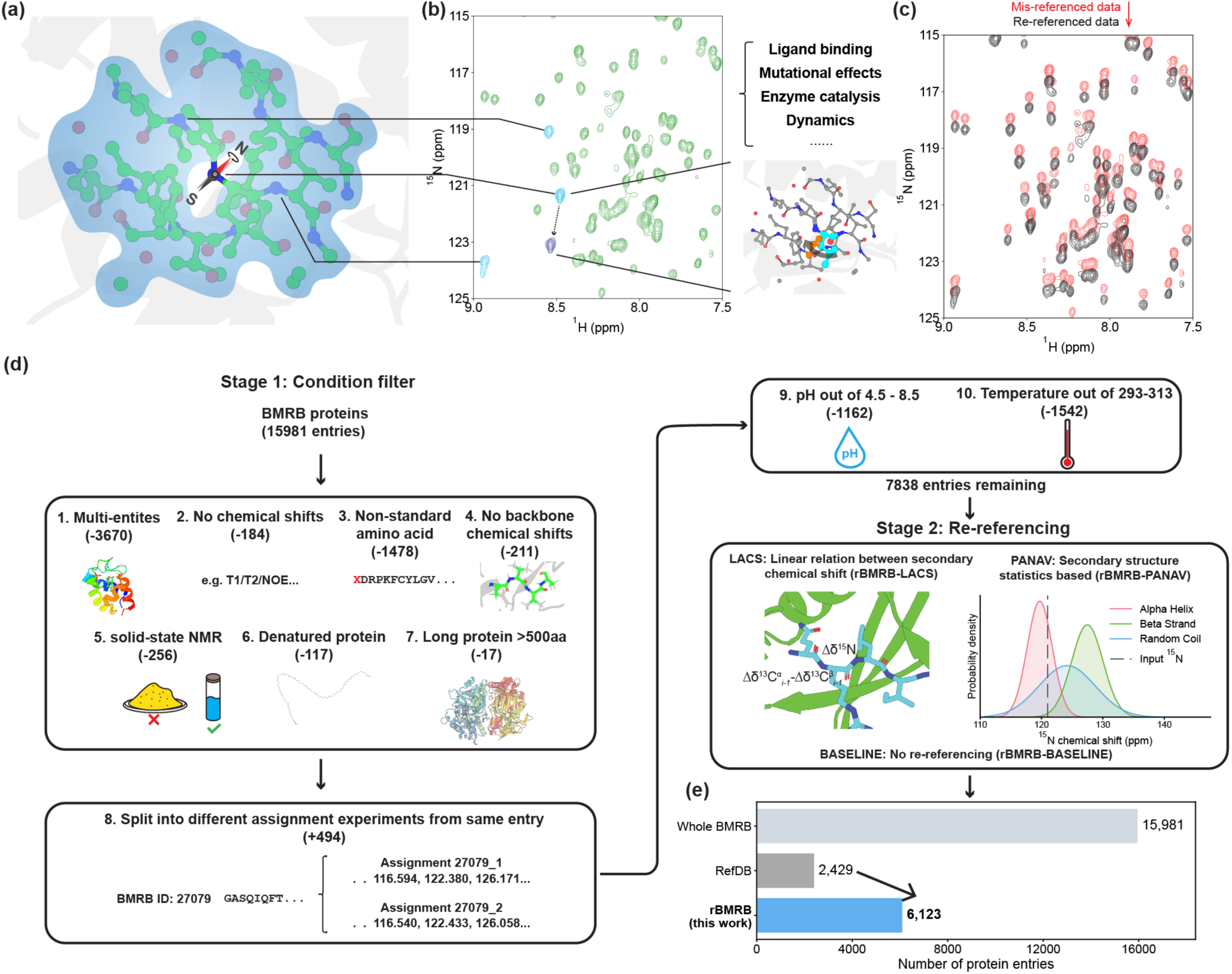
Curating a re-referenced dataset of all publicly available data (rBMRB) to assess chemical shift prediction. (a) Backbone chemical shifts report on the local chemical environment surrounding each atom. (b) Each peak in an ^1^H-^15^N HSQC spectrum reflects the local chemical environment of the corresponding backbone atom, making chemical shifts sensitive reporters of environmental perturbations. (c) Incorrectly referenced depositions carry a global offset of unknown magnitude across all chemical shifts (red arrow). Re-referencing methods estimate this offset to calibrate mis-referenced data (red) toward correctly referenced values (grey). (d) Data curation of rBMRB involved condition filtering (stage 1) and independent re-referencing via LACS^31,30^, PANAV^29,32^, and BASELINE (stage 2). This generated three datasets: rBMRB-LACS, rBMRB-PANAV and rBMRB-BASELINE (e) The curated rBMRB dataset retains 38% of the entries in the BMRB, substantially expanding on RefDB^25^.

Recently developed chemical shift predictors leverage learned representations of local chemical environments^5,6^ and report state-of-the-art performance. However, they remain within the same error magnitude as older models trained on just hundreds of proteins like SPARTA+ and SHIFTX2. Particularly for backbone ^15^N shifts^5,6,16,20–23^, prediction errors are large relative to the typical range of chemical shift values (Fig. S1, Table S1), making ^15^N predictors unusable for technical advancements in NMR data processing and prompting us to ask what these models may be failing to learn.

A natural resource for evaluating model performance is the Biological Magnetic Resonance Bank (BMRB), the largest repository of NMR data (>17,000 entries)^24^. However, one significant limitation of the BMRB is inconsistency in deposited datasets due to chemical shift mis-referencing^25^. Because chemical shifts are reference-dependent, differing conventions yield systematically different values for the same measurement, injecting systematic noise which is possibly harmful to model training^26^ (Fig. 1c). The main existing corrected database, RefDB, re-references entries using structure-based chemical shift prediction^25^ and is the source of training data for several predictors^5,27^. However, RefDB curation does not explicitly filter out confounding factors such as intermolecular interactions or atypical experimental conditions, potentially compromising its utility and reliability for downstream applications, including training monomer-based prediction models that assume negligible effects from these factors^5,8,27,28^.

To enable a fair evaluation of how well structure-based models capture local chemical environments, we curated a new dataset from the BMRB, which we call “rBMRB”. We applied many screening criteria on BMRB to retain only chemical shift entries of soluble, monomeric proteins, and performed chemical shift re-referencing using two probabilistic structure-free methods^29–32^.

Using rBMRB, we evaluated the effects of re-referencing on chemical shift prediction basing our analyses on custom chemical shift predictors fine-tuned based on the SLAE structure encoder (SLAE-CS)^6^. To our surprise, we found that re-referencing did not perceptibly change model performance. In that light, we trained models with substantially more data and found that more data also did not improve model performance. We analyzed essentially duplicate entries to estimate a median intrinsic accuracy limit of roughly 0.4-0.5 ppm for chemical shift prediction, yet the best models achieving median of 0.9 ppm are still far above this.

We next asked whether specific, predefined structural contexts such as solvent accessibility and secondary structures are systematically associated with higher or lower prediction error. We tested these differences for statistical significance across predictors built on four different foundation model representations – SLAE^6^, MACE-OFF23^33^, ESM3^1^ and ProteinMPNN^34^ – and validated the resulting patterns using microsecond molecular dynamics (MD) simulations. We found that the characteristic dynamics of exposed loop residues makes their chemical shifts easier to predict. Furthermore, increased prediction error is associated with atypical chemical environments, such as β-branched residues on loops and the combined effects of local packing and dynamics, some of which may be associated with specific protein functions.

Given these common themes we could identify, we were curious: could a structure-based model learn to anticipate its own errors directly from structure? We trained an error predictor (SLAE-ERR) using the top-performing model, SLAE-CS, taking its prediction errors as training targets, and achieved a net mean absolute error (MAE) of 0.760 ppm on error prediction, which is about two-fold larger than MAE of an oracle error predictor based on intrinsic uncertainty estimate. Our investigation of the discrepancies in SLAE-ERR underscored that places where it is most inaccurate are regions with extensive dynamics after excluding data quality issue. Beyond chemical shift uncertainty estimation, this learned error serves as a probe for model interpretability, identifying how protein foundation models capture local chemical environments and providing a more informative assessment of model confidence.

## Results

### Careful curation recovers a substantially larger usable dataset than prior resources

NMR spectroscopy enables the study of macromolecules in solution, and depositions in the BMRB span a wide range of molecular contexts (e.g. complexes, bound cofactors, ions) and experimental conditions (e.g. denaturation conditions, pH and temperature). Both contexts introduce variability in observed chemical shifts. Therefore, to study the effect of re-referencing on data, we needed to curate a new dataset to isolate re-referencing effects from other sources of chemical-shift variability (Fig. 1d; Methods).

Starting from 15,981 BMRB protein entries, we filtered to apo monomeric proteins in non-extreme conditions, removing more than half the entries and yielding 7,838 curated entries (Fig. 1d; Methods). The two largest sources of exclusions were entries containing multiple entities (e.g. bound metal ions, nucleic acids, proteins, etc.; removed 3,670 entries) and entries characterized outside a defined pH or temperature range (removed 2,704 entries; pH and temperature ranges: 4.5-8.5 and 293-313 K). The remaining filtered entries reflected structural or experimental incompatibilities, such as proteins with non-standard amino acids, denatured proteins, and chemical shifts from solid-state NMR experiments.

We then applied two types of probabilistic chemical shift re-referencing to the remaining entries to further reduce mis-referencing: PANAV^29,32^ and LACS^31,30^. Re-referencing was performed with the Python implementations of each method available in the package “makeshift”^35^, along with a no-re-referencing option (BASELINE). Our final dataset contained 6,123 entries, more than twice the size of RefDB (Fig. 1e). To our knowledge, this “rBMRB” dataset is the largest curated and rigorously filtered resource of protein chemical shift datasets.

### Re-referencing does not improve ^15^N chemical shift prediction accuracy

With these datasets in hand, we tested whether training SLAE-CS^6^, a state-of-the-art chemical shift predictor based on the SLAE protein structure encoder, on re-referenced data improves prediction performance (Fig. 2a; Methods). We focused on ^15^N chemical shifts, as these are among the worst-predicted chemical shifts^5,6,20,22^ and more accurate ^15^N prediction would provide improvements for NMR data processing. The overall ^15^N chemical shift distributions across our three paired datasets (rBMRB-BASELINE, rBMRB-PANAV, and rBMRB-LACS) were nearly identical (Fig. S2). rBMRB-LACS showed the largest average offset of 0.54 ppm from rBMRB-BASELINE, whereas the average offset for PANAV re-referencing was -0.02 ppm (Fig. 2b).

**Fig. 2.**
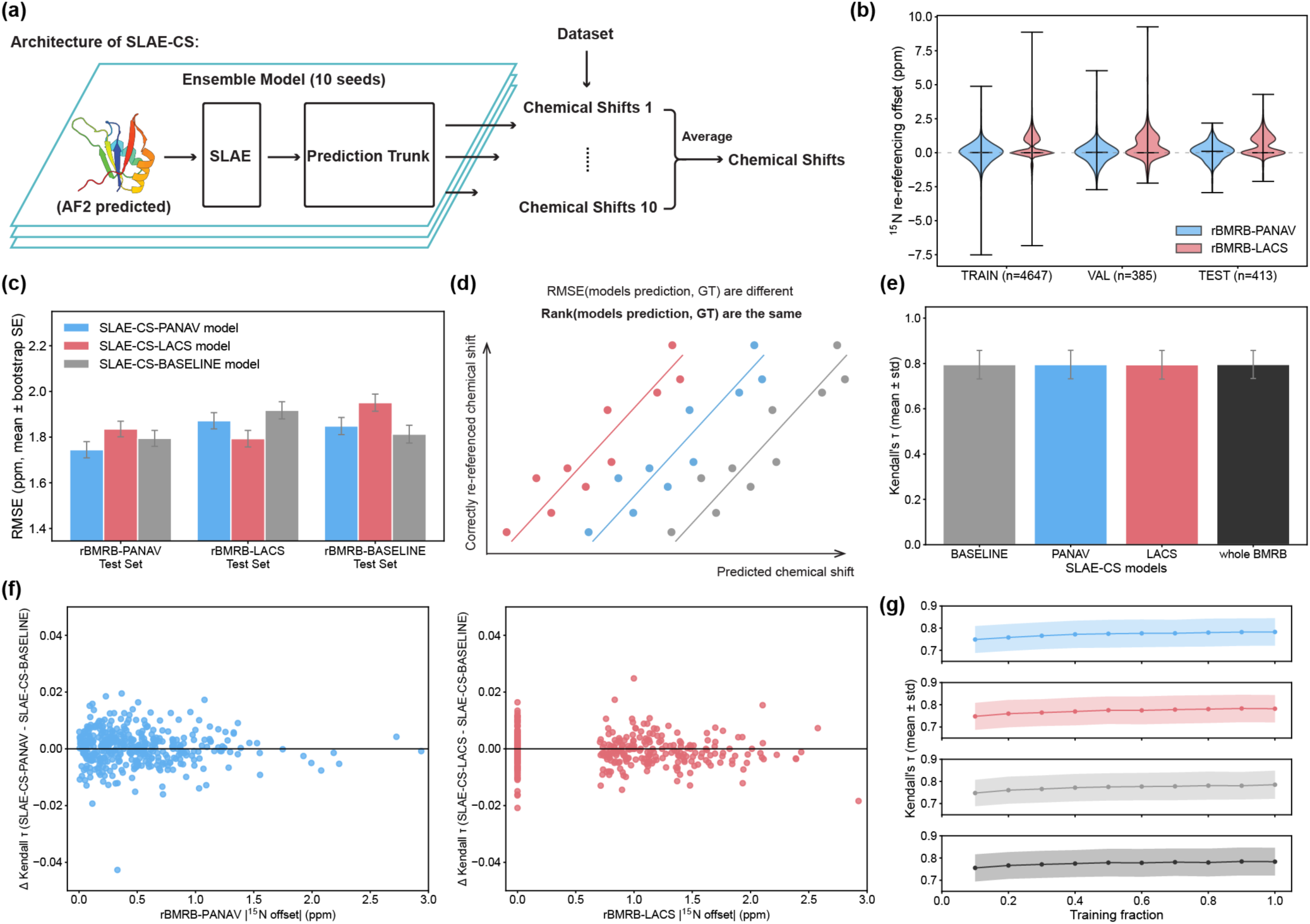
Re-referencing does not improve backbone ^15^N chemical shift prediction. (a) We tested the effect of re-referencing in the following chemical shift prediction architecture: an ensemble of ten independently trained models (AF2-predicted input structures, encoding via SLAE, followed by prediction trunk) generates chemical shift predictions that are averaged to obtain the final ensemble prediction. (b) Distributions of per-deposition ^15^N re-referencing offsets for rBMRB-PANAV and rBMRB-LACS (rBMRB-PANAV, μ = -0.02, σ = 0.73 ppm; rBMRB -LACS, μ = 0.54, σ = 0.87 ppm). Offset distributions are consistent across all train, val and test set splits. Horizontal lines indicate data median. (c) Test-set RMSE (mean ± bootstrap s.e.) for models trained on each re-referencing dataset – SLAE-CS-PANAV (blue), SLAE-CS-LACS (red), SLAE-CS-BASELINE (gray) – evaluated against each test set (SLAE-CS models -PANAV / -LACS / -BASELINE: rBMRB-PANAV test set 1.745 / 1.835 / 1.795; rBMRB-LACS test set 1.872 / 1.793 / 1.917; rBMRB-BASELINE test set 1.848 / 1.951 / 1.813; bootstrap SE ≈ 0.035). (d) Schematic illustrating relationship between rank-based metrics and referencing offsets. A global reference offset shifts predicted chemical shifts and therefore RMSE relative to ground-truth (GT) but preserves the rank order. (e) Kendall’s τ (mean ± s.d.) evaluated on the shared rBMRB-PANAV test set is nearly identical for SLAE-CS-PANAV, SLAE-CS-LACS, SLAE-CS-BASELINE (training set size = 4,647). Retraining on the complete, unfiltered BMRB (whole BMRB; training set size = 12,054) yields only a marginal improvement in Kendall’s τ (SLAE-CS models: -BASELINE / -PANAV / -LACS / -whole BMRB: 0.7948, 0.7953, 0.7943, 0.7958; standard deviation ≈ 0.063). (f) Per-protein change in Kendall’s τ relative to the SLAE-CS-BASELINE model, plotted against the magnitude of each protein’s ^15^N re-referencing offset, for models trained on rBMRB-PANAV (left, blue) and rBMRB-LACS (right, red). Gains and losses in ranking performance are of comparable magnitude and frequency with no systematic dependence on offset size (PANAV: r = −0.055, 95% CI [−0.140, 0.030], Pearson p = 0.27; LACS: r = -0.106, 95% CI [−0.201, -0.007], Pearson p = 0.033). (g) Mean Kendall’s τ as a function of training-set fraction for the SLAE-CS-PANAV (blue), SLAE-CS-LACS (red), SLAE-CS-BASELINE (ligh gray), SLAE-whole BMRB (dark gray) models (shaded band, ± s.d. across proteins). Performance improves most at small fractions and increases only marginally thereafter.

We partitioned each dataset into training, validation, and test sets splitting on 30% sequence similarity and structure similarity of 0.3 TM-score (training/validation/test set split: 4647/385/413), yielding comparable offset distributions across all splits (see Methods, Fig. 2b). Evaluation of the three models (SLAE-CS-PANAV, SLAE-CS-LACS, SLAE-CS-BASELINE) illustrated that each model performed best on chemical shifts originating from the same re-referencing method it was trained on; that is, SLAE-CS-PANAV performed best if the test set was re-referenced with PANAV whereas SLAE-CS-LACS performed best if it was re-referenced with LACS (Fig. 2c). This suggested that RMSE is insufficient to determine whether re-referencing improves prediction, as differences in RMSE can reflect systematic offsets introduced by re-referencing rather than differences in predictive performance.

To more robustly assess model performance to referencing artefacts, we evaluated Kendall’s τ as a per-protein rank-correlation coefficient: re-referencing introduces a systematic offset that biases RMSE but leaves prediction rank order unchanged (Fig. 2d). All three models yielded statistically indistinguishable Kendall’s τ distributions (Fig. 2e), indicating that re-referencing does not improve predictor performance. By decomposing Kendall’s τ at the protein level, we found no correlation between a predictor’s relative performance to the baseline and the magnitude of its corresponding re-referencing offset (Fig. 2f, Pearson/Spearman r = −0.055, p = 0.27 for PANAV; r = -0.106, p = 0.033 for LACS; bootstrap 95% CIs: [−0.140, 0.030] and [−0.201, -0.007]), further underscoring that re-referenced data did not improve predictive power.

Since re-referencing offered little benefit, we next asked whether simply adding more (unfiltered) training data could help. We trained models on the complete, unfiltered BMRB dataset of 12,054 entries (previously regarded to be too noisy for direct training^20,25^) and evaluated against the same test set. Kendall’s τ improved only modestly (∼0.002) despite the near doubling of training data (Fig. 2e), indicating that while the model can extract signal from a noisy dataset, model performance was already nearly saturated on the smaller, filtered datasets. Training on increasing fractions of the rBMRB (-PANAV, -LACS, -BASELINE) and the whole BMRB reaffirmed this hypothesis: performance improved sharply at small fractions, followed by only marginal gains thereafter (Fig. 2g). Notably, performance with 60% of the training data achieved a mean Kendall’s τ similar to that obtained with the full dataset (e.g. 0.7764 to 0.7827 for SLAE-CS-PANAV). Similar scaling trends have been previously observed when training protein structure prediction models^36^.

Given results in this section, and unless otherwise noted, we proceeded with only using the SLAE-CS-PANAV model and rBMRB-PANAV test set in all subsequent analyses because of its stricter outlier flagging mechanism (see Methods).

### Chemical shift prediction approaches an intrinsic noise floor set by BMRB itself

The previous analyses suggested to us that data quantity is likely not the limiting factor. We next asked whether the training data itself imposes an intrinsic accuracy limit^37,38^. To estimate the extent to which variability in deposited chemical shifts may limit prediction accuracy, we quantified disagreement between sequence-identical depositions collected under closely matched experimental conditions as an accuracy ceiling. We first clustered the dataset at 100% sequence identity after Stage 1 condition filtering (Fig. 1d). Next, we retained only pairs whose temperature and pH differed by <2.5 K and <0.2 units, respectively, to isolate irreducible noise from condition-dependent effects (Fig. S6). Pairs with poorly documented metadata but substantially different chemical environments were manually inspected and excluded for major differences in binding, oligomerization state, redox state, denaturing conditions, post-translational modifications, or membrane environment. This retained pairs of depositions that differed only in minor experimental details, such as sample concentration and buffer composition, yielding 72 sequence-identical pairs (Methods, Table S2). Table S3 provides details of all pairs manually excluded due to ambiguous metadata documentation. The screening process could be further refined if additional metadata were available and incorporated into the chemical shift prediction model.

For each pair, we decomposed the raw disagreement into a referencing offset, estimated as described above, and an intrinsic, reference-independent noise term (Fig. 3a, Methods). LACS and PANAV re-referencing gave comparable results. Pooled across pairs, the intrinsic RMSE substantially exceeded the referencing offset by approximately three-to four-fold (1.56 vs. 0.40 ppm for PANAV and 1.91 vs. 0.64 ppm for LACS; Fig. 3b and Fig. S7), so correcting referencing offsets removes only part of the discrepancy. This pooled intrinsic RMSE is comparable to the ^15^N prediction RMSEs reported above (1.75 ppm for SLAE-CS-PANAV and 1.79 ppm for SLAE-CS-LACS), but notably the per-pair distribution is heavy-tailed (Fig. 3c), with a much lower median of 0.48 ppm. This phenomenon mirrors the broader, outlier-dominated distribution of chemical shift prediction errors (Fig. S5). Omitting our manual metadata review changes the median modestly (0.58 vs. 0.48 ppm) but added pairs to the high-RMSE tail, so careful metadata curation matters more for outliers rather than the typical pair (Fig. 3c).

**Fig. 3.**
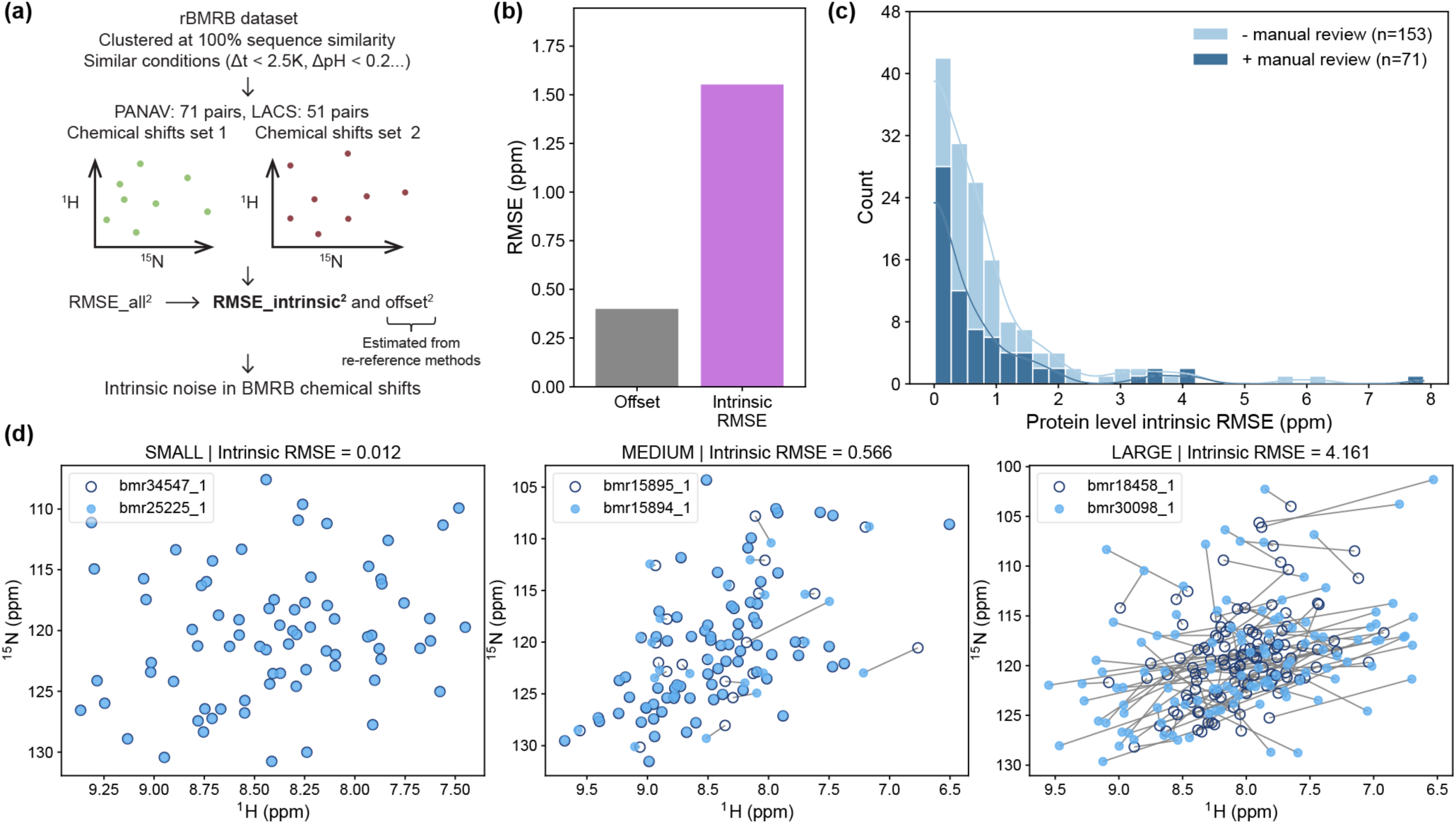
Estimating the BMRB’s intrinsic noise floor for chemical shift prediction. (a) Approach for estimating the intrinsic accuracy ceiling of chemical shift prediction. Within the rBMRB dataset, we identified 72 pairs of depositions sharing 100% sequence identity under matched experimental conditions. The total disagreement between each pair’s depositions’ chemical shifts is decomposed into RMSE_intrinsic^2^ and offset^2^ to separate an irreducible intrinsic noise term from a global referencing offset. (b) Aggregate referencing offset and intrinsic RMSE (pooled root mean square) by referencing method PANAV (n = 71). The intrinsic noise (1.56 ppm) is roughly four-fold larger than the corresponding referencing offset (0.40 ppm), indicating that most of the disagreement between sequence-identical depositions likely reflects irreducible noise for predictors. (c) Distribution of per-pair intrinsic RMSE for PANAV before and after last round of manual review (before: light blue, n = 153, median = 0.58; after: dark blue, n = 71, median = 0.48 ppm). After manual review most pairs show intrinsic RMSE <2 ppm, with a heavy tail at >3 ppm. Curves are kernel density estimates. (d) Representative offset-corrected (by PANAV) pairs spanning the small (left), medium (middle), and large (right) relative intrinsic RMSE shown as overlaid ^1^H-^15^N HSQC spectra, where lines connect matched residue pairs.

We illustrate representative pairs at a variety of intrinsic RMSE values with simulated ^1^H-^15^N HSQC spectra in Fig. 3d. As would be expected for identical constructs under identical conditions, we show in (a) human secreted Ly-6/uPAR-related protein 1 (entries bmr25225_1 vs. bmr34547_1^39^ at pH 4.7 and 310 K) with near-zero RMSE after offset correction. In (b) we display two spectra of periplasmic protein PCu_A_C (entries bmr15894_1 vs. bmr15895_1 at pH 7.2 and 298 K) with modest shifts among a subset of peaks. This can be explained by existence of two conformers^40^. However, in (c) are two depositions with substantial per-residue disagreement, a 122-residue HIRAN domain of human HLTF (entries bmr18458_1 vs. bmr30098_1^41^, at pH 6.8 and 298 K). Although both entries were deposited by the same research group, we could not determine a source of discrepancy.

### Protein foundation models struggle to capture atypical chemical environments

We next asked whether specific structural features are systematically associated with poor prediction performance. Using a common pipeline (Fig. 4a), we evaluated five foundation models ranging from structure only (ProteinMPNN) to sequence only (ESM3) to assess the extent to which each representation captures local chemical environments. SLAE-CS performed the best at chemical shift prediction overall, followed by ESM3 with sequence and structure input and MACE-OFF23. These three most accurate predictors showed the highest agreement with each other (Pearson correlation = 0.72-0.80; Fig. 4b-c).

**Fig 4.**
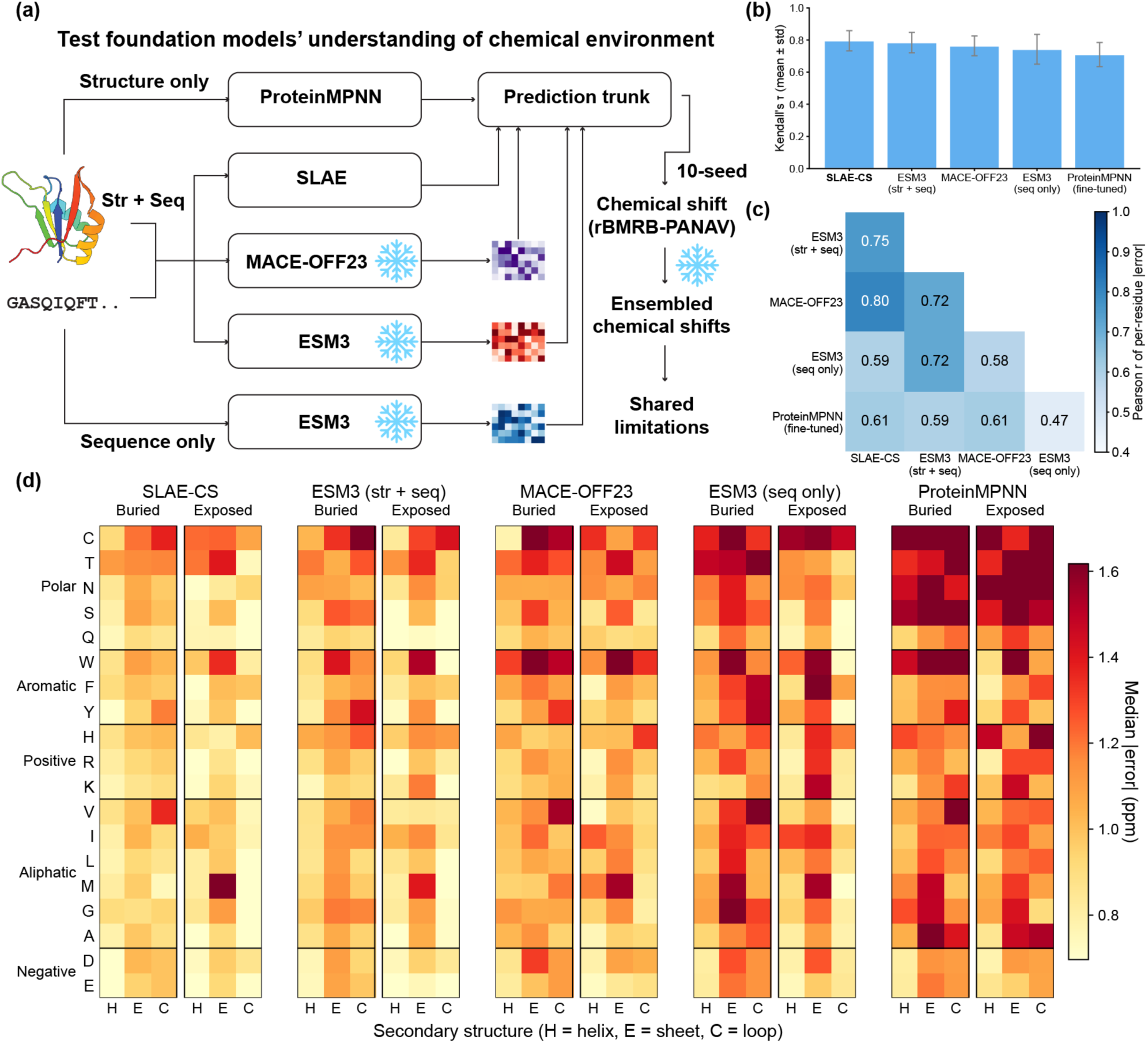
Protein foundation models share residue- and context-specific chemical shift prediction errors. (a) Schematic of the workflow for evaluating foundation model representations. Protein structure and/or sequence inputs are encoded by ProteinMPNN (structure only), SLAE, MACE-OFF23 or ESM3 (structure + sequence, or sequence only). The resulting embeddings are passed to a shared prediction trunk to predict chemical shifts based on rBMRB-PANAV and get ensemble-averaged chemical shifts for downstream evaluation (snowflake indicates frozen model). (b) Kendall’s τ between predicted and experimental chemical shifts (mean ± s.d. across proteins) for each predictor (SLAE-CS, ESM3 (str + seq), MACE-OFF23, ESM3 (seq only), ProteinMPNN (fine-tuned): 0.80, 0.78, 0.76, 0.74, 0.71). (c) Agreement in per-residue ^15^N absolute prediction error between every predictor, measured by Pearson r. (d) Median ^15^N absolute prediction error stratified by residue type, secondary structure (H, helix; E, strand; C, loop) and solvent accessibility (buried, exposed) for each predictor. Residue types are grouped as polar, aromatic, positively charged, aliphatic and negatively charged.

We examined how prediction errors are associated with secondary structure^42^, solvent accessibility^43^, and residue identity^44^, and found that errors fall mostly between ∼0.7 and ∼1.6 ppm, with some consistently poorly predicted classes, such as Trp and Met on exposed β strand, shared between SLAE-CS and other predictors (Fig. 4d). We proceeded with using SLAE-CS for subsequent analyses. Intriguingly, the top-performing models in Figure 4b differ in some categories (i.e., buried valines in loops), suggesting avenues for further exploration and model interpretation.

Stratifying prediction error by secondary structure and solvent exposure, we found that exposed loop residues were predicted more accurately than residues in any other category (Fig. 5a, Fig. S8). We hypothesized that this reflects conformational averaging on fast timescales, which would smooth the ensemble-averaged shift toward a value that is easier to predict (Fig. S9). To test this, we ran 1 µs molecular dynamics simulations of 48 proteins from the test set, selected to span a range of lengths and of α-helical and β-strand content (Fig. 5b, see Methods). Indeed, among exposed loop residues, proteins with greater backbone dihedral circular variance had lower ¹⁵N chemical shift prediction error [ρ = -0.58, p = 1.6e-5] (Fig. 5c), consistent with this hypothesis (Fig. S10).

**Fig. 5.**
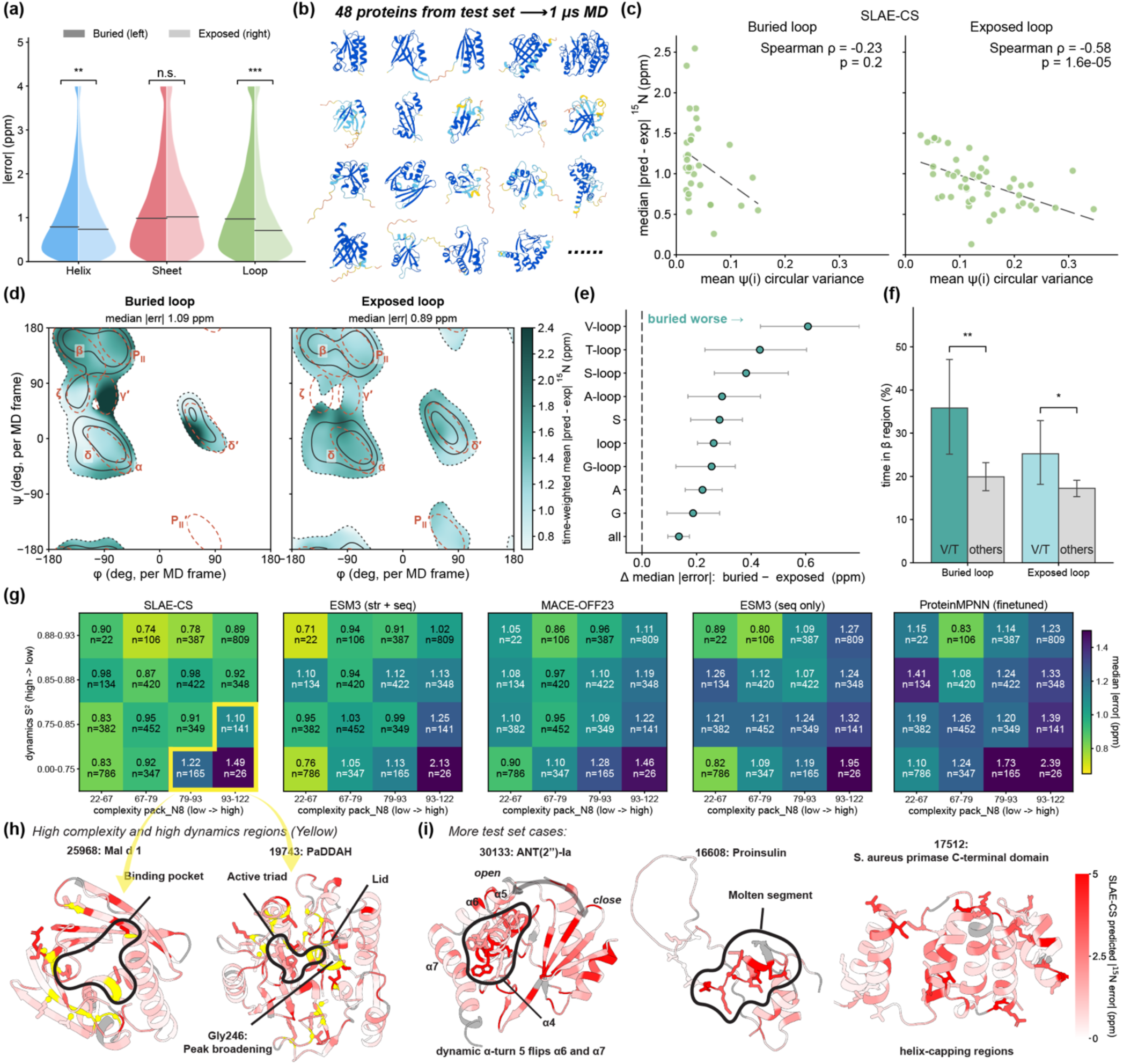
Atypical backbone conformations, dynamics and packing underlie high chemical shift prediction errors, often associated with functional sites. (a) ^15^N SLAE-CS |error| categorized by secondary structure. Each violin is split into buried residues (left half) and exposed residues (right half). Horizontal lines mark medians. Brackets compare buried with exposed residues (**, p < 0.01; ***, p < 0.001; n.s., not significant). (b) One-microsecond MD simulations of 48 SLAE-CS test-set proteins were used to investigate patterns in chemical shift errors. Nineteen representative proteins are shown. All 48 proteins feature mixed secondary structures and high-confidence AlphaFold2 starting structures (see Methods). Structures are colored by pLDDT (dark blue, >90; light blue, 70–90; yellow, 50–70; orange, <50). (c) Per-protein median ^15^N |error| of SLAE-CS against the mean MD circular variance of ψ(i), for buried (left) and exposed (right) loop residues. Each point is one protein (proteins with ≥ 5 residues in that class). Dashed lines are linear fits; Spearman ρ and p are shown in each panel. (d) Per-frame MD Ramachandran maps of buried (left, n = 350) and exposed (right, n = 871) loop residues. Black contours enclose 50%, 80% and 95% of frames (dotted line, 95%), with every residue weighted equally. Shading is the time-weighted mean SLAE-CS ^15^N |error| of the residues visiting each (φ, ψ) bin. Dashed outlines are the Ramachandran regions of Hollingsworth & Karplus (2010)^45^. (e) Difference in median absolute prediction error from SLAE-CS between buried and exposed residues for the full test set stratified by secondary structure, residue type, and residue–secondary structure combinations. Error bars indicate 95% confidence intervals from 100,000 protein-cluster bootstrap resamples; significance assessed with Bonferroni correction for family-wise absolute error across all tested strata (strata shown are those significant at p < 0.01). (f) Mean fraction of MD time spent in the β region by Val/Thr compared with all other residues, in buried and exposed loop. Error bars are 95% CIs from 5,000 bootstrap resamples over proteins; brackets give the two-sided bootstrap p for the difference (*, p < 0.05; **, p < 0.01). (g) Median ^15^N absolute prediction error binned jointly by local packing complexity (pack_N8 quartiles, number of heavy atoms within an 8 Å radius of N) and backbone order parameter S^2^ (quartiles). n, residue counts per cell. Yellow outlines mark the high-complexity, low-S² regime examined in (h). (h) Representative SLAE-CS case studies from the regime outlined in (g): the apo-state structure of Mal d 1 (BMRB 25968; left) and the apo-state structure of PaDDAH (BMRB 19743; right). Structures are colored by SLAE-CS-predicted ^15^N absolute error (white, low; red, high; grey, no data). Residues in high-complexity, high-dynamics regime are highlighted in yellow and shown as sticks. In PaDDAH, the active triad and Gly246 are also shown as sticks. (i) Additional representative test-set cases: the apo-state structure of ANT(2″)-Ia (BMRB 30133; left), proinsulin (BMRB 16608; middle) and the C-terminal domain of S. aureus primase (BMRB 17512; right). Coloring is as in (h). Atypical regions associated with conformational change, biological function or non-canonical geometry are shown as sticks.

We next investigated why buried loop residues are predicted less accurately than exposed ones. Ramachandran distributions showed that buried loop residues populate the β-strand region more than exposed loop residues do (Fig. 5d). Buried loop residues also exhibited a broader distribution of dihedral angles near the ζ region^45^, indicating greater diversity in the backbone conformations (Fig. 5d). Higher prediction errors were primarily concentrated in β and P_II_ regions (Fig. S11).

We examined solvent-accessibility-associated differences across more fine-grained structural categories (see Methods) and noticed that β-branched residues valine and threonine showed the biggest differences in error between buried and exposed loops (Fig. 5e, Fig. S8). We also observed that both residues showed higher occupancy in β-region of the Ramachandran plot, regardless of solvent-exposure state, which may account for the larger differences in prediction error (Fig. 5f).

Together these findings demonstrate that models perform worse on loop residues that adopt atypical conformations, particularly for buried loop residues trapped in β-regions, compared with other contexts. We created a metric that allows us to simply summarize these effects across all residues. Categorizing residues by their MD-derived backbone order parameter (S^2^) and the number of heavy atoms within an 8 Å radius of N (pack_N8) shows that highest error is present in residues with low S^2^ and high pack_N8 (Fig. 5g).

What are these residues with high packing density and low S²? In investigating, we found this type of prediction difficulty was often associated with biologically-relevant dynamics. In apo Mal d 1 (BMRB 25968, Fig. 5h), several binding-pocket residues with high error are known to undergo micro-to-millisecond motions^46–48^. In apo DM-PaDDAH (BMRB 19743; Fig. 5h), high-error residues cluster in one contiguous region spanning the active-site lid and catalytic triad. This lid contains picosecond-to-nanosecond motion related to function in relaxation experiments^49^.

Other poorly predicted proteins in the test set involve atypical chemical environments that may be difficult for protein foundation models to learn (Fig. 5i). In apo ANT(2″)-Ia (BMRB 30133), α-turn 5 has been captured in both open and closed conformations found in the PDB, suggesting that the aminoglycoside-binding α4 pocket is gated by conformational dynamics^50^. Proinsulin’s “molten segment” (BMRB 16608) is poorly predicted and exhibits flexibility at timescales slower than overall tumbling^51^. In the C-terminal domain of *S. aureus* primase (BMRB 17512), errors concentrate at helix-capping regions, consistent with their distinct hydrogen-bonding interactions^52^. More generally, helix capping residues show higher error than canonical helical residues across the test set (Fig. S12). Together, these examples demonstrate how evaluation with the rBMRB dataset can help identify specific structural features that predictors may fail to learn.

### A structure-based error model anticipates prediction difficulty, but is still hampered where dynamics exist

Beyond characterizing difficult-to-predict chemical environments, we were curious if a structure-based model could also learn to anticipate its own errors like pLDDT in AlphaFold2^53^. As a baseline, we assembled an explicit set of structural descriptors previously implicated in chemical shift prediction^20–22^ including backbone conformation, ring-current effect, structure reliability (pLDDT), local packing density, hydrogen-bonding geometry, and dynamics (Fig. 6a; see Methods). We framed this task as a binary classification problem: classify the hardest 10% versus easiest 50% of test-set residues by SLAE-CS’s absolute error (Fig. S13). A baseline classifier achieved an out-of-fold AUROC of 0.633 (Fig. 6b), and incorporating the full descriptor set raised the AUROC to 0.763. Permutation importance ranked local packing density and backbone order parameter (S^2^)^54^ as the most informative descriptor groups, followed by predicted μs–ms conformational exchange from Dyna-1^15^ and feature group describing backbone conformation including psi angles and Foldseek tokenization (Fig. 6c, Fig. S14; See Methods). This supports our previous characterization that S^2^ and local packing can describe chemical shift prediction errors, extended across a larger protein set (Fig. S15-16).

**Fig. 6.**
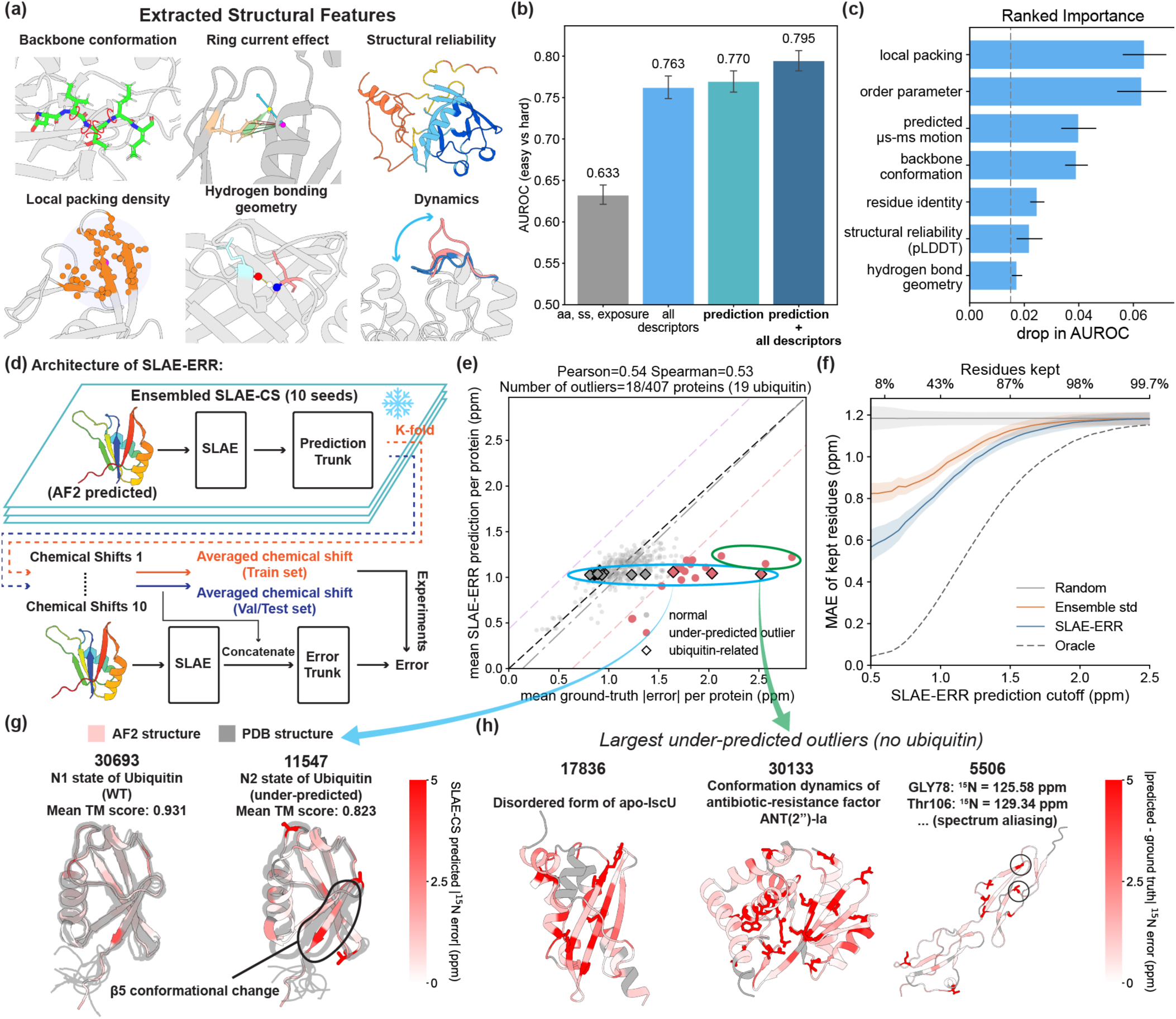
A structure-based error model SLAE-ERR captures overall prediction difficulty, but is fundamentally limited by the finite information a single static conformation can encode. (a) Explicit structural feature categories evaluated. (b) Out-of-fold AUROC for the hard-versus-easy classification task using the combined baseline with residue identity, secondary structure, and solvent accessibility (gray, AUROC: 0.633); the full set of explicit structural features from (a) (blue, AUROC: 0.763); SLAE-ERR’s predicted absolute error (green, AUROC: 0.770); or explicit structural features and SLAE-ERR’s predicted absolute error (navy, AUROC: 0.795). Error bars, s.d. across folds. (c) Group permutation importance for the hard-versus-easy classification task, quantified as the decrease in AUROC upon shuffling each feature group within each of the K cross-validation folds (see Methods). Error bars indicate the standard deviation across folds. Only groups with permutation importance larger than 0.015 are shown. The local packing density and backbone order parameter (S^2^) carry the most discriminative signal. (d) Architecture of the SLAE-ERR. An ensemble model of ten frozen SLAE-CS models trained on different seeds to predict backbone ^15^N chemical shifts from an AF2-predicted structure (snowflake indicates frozen model); the mean and standard deviation of the model ensembles are concatenated with the structural representations and passed as additional input to a separate error-prediction trunk. (e) Mean SLAE-ERR predicted versus ground-truth ^15^N absolute error per protein in the test set (n = 407). Grey dash-dot line, linear fit; black dashed line, y = x. Protein is an outlier if residual ≥ Q3 + 1.5 IQR (red, under-predicted; n = 18) or ≤ Q1 − 1.5 IQR (purple, over-predicted; n = 0). The two bounds are drawn as light dashed lines. Diamonds mark the 19 ubiquitin-related entries. Circled entries share nearly the same predicted error. (f) MAE of SLAE-CS ^15^N predictions for test-set residues retained at each SLAE-ERROR predicted-error cutoff. The top axis gives the fraction of residues retained. At each cutoff, the same number of residues was retained by lowest ensemble standard deviation (Ensemble std), at random (Random) or by lowest observed error (Oracle). Shaded areas, 95% ranges from 2,000 protein-cluster bootstrap resamples. (g) AF2 (pink) and PDB (grey) structures of ubiquitin in the WT N1 state (BMRB 30693) and the under-predicted N2 state (BMRB 11547), colored by SLAE-CS-predicted ^15^N absolute error (grey, no data). Mean TM-score between the AF2 model and the PDB models for states N1 (0.931) and N2 (0.823). (h) AF2 structures of the 3 non-ubiquitin entries with the largest error from (e), colored by |predicted − ground-truth| ^15^N error (grey, no data). BMRB entry 17836 is a disordered apo-IscU, and BMRB entry 30133 undergoes conformational dynamics not represented in the known PDB structure. In BMRB entry 5506, error is concentrated at residues with aliased ^15^N shifts, inflating the per-protein mean

We then trained an error prediction model using AF2-predicted structures, SLAE embeddings, and the mean and standard deviation of predictions from ten independently trained SLAE-CS models (Fig. 6d; see Methods). The resulting model, which we named SLAE-ERR, achieved a per-residue mean absolute error (MAE) of 0.760 and a root-mean-squared error of 1.228, with the oracle error predictor providing a lower bound of 0.35 MAE (see Methods). It also correlated with errors averaged at the protein level (Pearson’s r = 0.54, Spearman’s ρ = 0.53, n = 407; Fig. 6e) and modestly improved the binary classification task (Fig. 6b; AUROC of 0.770 with prediction alone and 0.795 when combined with the feature set). Filtering low-confidence predictions in the test set using SLAE-ERR predictions yielded a substantially lower MAE than filtering based on the SLAE-CS ensemble’s standard deviation alone or random sampling (Fig. 6f).

To understand remaining discrepancies in SLAE-ERR predictions, we examined protein outliers (Fig. 6e; see Methods). The rBMRB test set contains 19 ubiquitin entries, of which three have under-predicted errors (BMRB entries 18611, 11547, and 27512). Entries 18611 and 11547 are known to adopt alternative conformational states^55,56^, and entry 27512 exhibits spectral aliasing (Fig. S17), an incurable experimental artefact error. Across ubiquitin entries, the ground-truth absolute error ranges from 0.8 to 2.5 ppm, yet the predicted error stays close to 1.0 ppm.

The input structures for all entries are highly similar to wild-type (WT BMRB entry 30693; TM-score ≥ 0.985) despite sequence variation (≥90% identity; Fig. S18-S19). The model cannot distinguish between entries whose experimental conformational state differs from WT, likely because current structure-based methods are poor mutant effect predictors^57,58^ (Fig. S20). ESM3 with sequence-only input recapitulates this pattern (Fig. S21), suggesting the behavior is not specific to structure-based prediction. The AF2 model used by SLAE-CS and SLAE-ERR agrees with the WT structure (N1 state)^59^ and not the alternate state (N2 state) of the under-predicted BMRB entry 11547 in the C terminal where the structures deviate^55^ (Fig. 6g). Promisingly, when SLAE-CS is provided a correct structure model, the MAE in the C-terminal region decreases from 2.08 to 0.81 ppm (Fig. S22).

For other under-predicted outliers, we similarly observed that input structures did not match the relevant conformational state (Fig. 6h). For example, the apo form of IscU (BMRB 17836) is disordered^60^ unlike the AF2-predicted structure, and the antibiotic-resistance factor ANT(2”)-la (BMRB 30133) undergoes large-scale catalysis-associated conformational dynamics^50^. Spectral aliasing remains an artefactual problem: for the central complement control factor^61^ (BMRB 5506), the largest ground-truth error are concentrated at several residues which are well outside their typical ^15^N shift range (Gly73, 125.58 ppm; Thr106, 129.34 ppm, Table S1^24^) due to spectral aliasing. In summary, the largest disagreements between predicted errors from SLAE-ERR and ground-truth error arises primarily when the input structure fails to capture the chemical environment reflected in the experimental chemical shifts.

## Discussion

Deep learning has revolutionized our ability to predict protein structures and design proteins. However, there has been an increasing effort to improve their ability to predict and manipulate fine-grained chemical environments fundamental to biological function^62^. Furthermore, there is substantial need across many deep learning for biology contexts to better interpret what models are learning. By evaluating state-of-the-art deep learning models on chemical shift prediction, our results demonstrate that these foundation models struggle to capture atypical environmental contexts. These findings extend beyond secondary structure labels to encompass environments central to protein function (e.g. active sites, binding pockets), crucial geometries (e.g. helix-capping), and protein dynamics (e.g. alternate states, conformational changes), highlighting the rich biological insights that chemical shift prediction can provide. Protein foundation models have been previously shown to struggle with related tasks including mutant effect prediction^57,58^, pKa prediction^63^, and conformational ensemble prediction^64^. A similar asymmetry has also been observed in protein design, where diffusion-based backbone generative models are biased towards α-helices^65,66^. Together, these findings suggest a broader challenge for foundation models in representing structural context faithfully, particularly for chemistries depending on non-local interactions.

We show that some failure modes are predictable by these models: our model SLAE-ERR, trained to predict chemical shift errors, is able to flag atypical environments where chemical shift predictions may be unreliable without requiring experimental input. Such model calibration is routine for other tasks^53^, but it is not a common practice for chemical shift prediction^5^. SLAE-ERR achieved a MAE of 0.760 ppm. To contextualize this, an “oracle” error predictor assuming our estimated intrinsic noise floor of 0.5 ppm is able to achieve 0.35 ppm (see Methods). This indicates SLAE-ERR is roughly two-fold higher error than an oracle error predictor. Our results also suggest that chemical shift prediction error might be a way to guide other models to develop chemically interesting protein designs, for example as an indicator of geometry-or function-related complexity.

We provide the curated datasets herein for use by the structural biology and deep learning communities. The rBMRB provides re-referenced and curated datasets, along with condition-annotated chemical shifts. This provides a rich dataset for broader use beyond chemical shift prediction, such as chemical shift perturbations from ligand binding, mutant effects, or environmental conditions. Our hope is that these data may also be useful for finetuning models in other tasks of protein deep learning, such as mutant effect prediction, and model interpretability efforts.

Looking forward, many of the failure modes of current foundation model stem from features that cannot be captured by a single static structure, including conformational heterogeneity and biological response due to different conditions. As ensemble-averaged observables sensitive to these factors, chemical shifts provide a natural bridge between structure prediction and protein function. Continued curation of conformational ensembles, explicit measures of protein dynamics (e.g., relaxation-derived order parameters or exchange distributions), and experimental conditions under which these observables are measured could enable models to move beyond single-state structure prediction towards predicting protein motions in different environments. More broadly, chemical shifts and related NMR observables are poised to serve as benchmarks for evaluating model predictions and as training data and inference-time constraints, grounding protein foundation models in the physical ensembles that underlie biological function.

## Acknowledgments

The authors thank members of the Wayment-Steele lab, Katherine Henzler-Wildman, and Samuel Butcher for useful discussion. C.Z. was funded by E.W. Hopkins Biochemistry Graduate Fellowship administered by the University of Wisconsin-Madison Department of Biochemistry. C.Z. and H.K.W.S. acknowledge support from the University of Wisconsin - Madison Biochemistry department startup funds.

## Author contributions

C.Z., G. E. N., and H.K.W.S. designed the research. C.Z. performed data curation, trained models, ran molecular dynamics simulations, and analyzed data. C.Z. and Y.C. wrote code. C.Z., G. E. N., H.K.W.S. wrote the manuscript. All authors reviewed and edited manuscript.

## Data availability

The rBMRB chemical shift dataset, including the train, validation and test splits and AlphaFold2 structures, is available on Hugging Face (https://huggingface.co/datasets/hws-lab/rBMRB). Data underlying the figures, including molecular dynamics trajectories, and the inputs of the dataset curation are available on Zenodo (https://doi.org/10.5281/zenodo.24023681). Chemical shift data were obtained from the BMRB (https://bmrb.io).

## Code availability

Code for training, inference, figure analyses and dataset curation is available at https://github.com/WaymentSteeleLab/SLAE-CS-and-SLAE-ERR. Trained SLAE-CS and SLAE-ERR weights are available at https://huggingface.co/hws-lab/SLAE-CS-and-SLAE-ERR.

## Methods

### rBMRB curation

Before applying the structure-free re-referencing methods, we ensured that the dataset was as clean as possible so that the effects of re-referencing could be isolated from other unrelated sources of chemical-shift variability. Starting from the full BMRB (15,981 protein entries), we followed the filtering protocol shown in Figure 1d using the software package makeshift^35^.

In the first round, we removed all BMRB entries containing multiple entities (including binding partners such as metal ions, other proteins, nucleic acids, or ligands). These species can influence protein chemical shifts through direct or allosteric interactions. This step eliminated 3,670 entries, the largest single source of removal. The same rationale of restricting the dataset to structural contexts current frameworks can represent was applied to exclude entries containing non-standard amino acids (1,478 entries), ensuring that all structures could be generated using AlphaFold2^53^. Since structure prediction models tend to predict the native folded conformation regardless of denaturing experimental conditions, denatured proteins (117 entries) were excluded to avoid providing systematically incorrect structural inputs.

Additional filtering criteria targeted the specific requirements of backbone chemical shift prediction. Entries containing only relaxation experiment data (e.g., T1/T2/NOE) and lacking chemical shift assignments were removed (184 entries), as were entries containing only side-chain chemical shifts without backbone assignments (211 entries). Solid-state NMR entries were discarded (256 entries), and proteins longer than 500 residues were excluded (17 entries).

Finally, and the second-largest source of removal after the multi-entity filter, entries falling outside a defined pH (4.5–8.5) or temperature (293–313 K) window were removed (1,162 and 1,542 entries, respectively). Both parameters are known to systematically perturb chemical shifts, with pH acting through altered protonation states of titratable side chains and temperature through changes in thermodynamic averaging over conformational substates. They are difficult for a structure-only-based predictor to capture and that fall outside the scope of this study^28,67^.

Entries containing multiple assignment experiments from a single BMRB entry were split into separate entries to maximize data utilization and improve training robustness, as such experiments typically share the same protein sequence but yield distinct chemical shift values reflecting different experimental preparations (494 entries added). After these filtering steps, 7,838 BMRB entries remained as the curated dataset for subsequent re-referencing (Fig. 1d).

### Whole BMRB curation

To test the effects of more data, the whole BMRB was also used for training chemical shift predictor, regardless of the noise inside. Some necessary filtering was still applied specifically for backbone chemical shift prediction training, including removal of 1,010 entries without proteins inside, 911 entries without available backbone chemical shifts, 1859 entries with non-standard amino acids, and sequences longer than 500 residues. This resulted in 14,966 entries.

### Re-referencing Methods

Two structure-free re-referencing methods were used in this study: PANAV and LACS^29–32^. Although the analyses in this paper focus on backbone ^15^N shifts, both the condition-based filtering described above and the re-referencing procedure described here were applied jointly across all available backbone chemical shifts for each entry: ^15^N, ^1^H, ^13^C^α^, ^13^C^β^, and carbonyl ^13^C.

PANAV first infers the secondary structure of each residue from its chemical shifts, relying primarily on H^α^ shifts, which are largely insensitive to re-referencing errors (Fig. 1d). Using statistics of each atom type’s chemical shift distribution within different secondary structure contexts, PANAV assigns a secondary structure to each residue, then averages the differences between the statistical values and the observed shifts to obtain offsets for re-referencing^68^. In addition to these offsets, PANAV flags outliers and potential typographical errors; we applied these flags across all datasets’ validation and test sets to remove low-quality data points for more fair comparison. LACS exploits linear relationships between the differences in secondary chemical shifts of the preceding/current residue’s ^13^C^α^ and ^13^C^β^ and the secondary chemical shift of the atom type being re-referenced, since referencing errors should cancel out when computing differences of secondary chemical shifts.

Both methods were implemented using the “makeshift” Python package. Because PANAV’s flags are data-dependent, we additionally applied a general outlier filter to both the LACS and BASELINE datasets, removing data points that deviated from the BMRB statistics (Table. S1, fetched from BMRB on 4/17/2026 and used in makeshift as well) by more than five standard deviations. Note that all rBMRB-PANAV, rBMRB-LACS, rBMRB-BASELINE and whole BMRB used PANAV flags in their validation and test sets.

### Train/val/test set splits

To prevent structural leakage between data splits, protein structures were clustered prior to splitting. Structures corresponding to the 6,123 entries common to re-referencing related datasets (PANAV, LACS, BASELINE) were pooled and clustered using Foldseek easy-cluster^69^, with structural alignment via TMalign^70^ (--alignment-type 1), a TM-score threshold of 0.3, a minimum pairwise sequence identity of 30%, and bidirectional coverage (--cov-mode 0), yielding 4,087 clusters.

Clusters were randomly assigned (seed = 42) to training, validation, or test sets and all entries within a given cluster were placed together in the same split to prevent structural overlap across sets. This procedure yielded 5,304 training, 406 validation, and 413 test entries. Because cluster-level partitioning alone does not guarantee the absence of cross-split homologs, we applied an additional leakage filter while keeping the test set fixed. Homology was defined from a single exhaustive all-against-all MMseqs2^71^ search over all 6,123 candidate entries. Two entries were considered homologous if their alignment had ≥30% sequence identity, covered ≥80% of either sequence, and had E ≤ 10^-3^. Validation entries homologous to any test entry were removed, and training entries homologous to any test entry or to any retained validation entry were removed. Entries that shared a BMRB accession with an entry in another split were also removed. The final dataset contained 4,647 training, 385 validation, and 413 test entries. The same entry-level split was applied identically to all re-referencing related datasets, ensuring that all downstream comparisons used matched train/validation/test partitions.

For the whole BMRB, the validation (385 entries) and test (413 entries) sets were fixed to those used for rBMRB-PANAV, rBMRB-LACS and rBMRB-BASELINE, and a leakage-free training set was drawn from the remaining pool of 14,966 whole BMRB entries. Leakage was assessed on the combined set of pool, validation and test entries in two steps. First, all structures were clustered with Foldseek easy-cluster using the same parameters as above (TMalign alignment, TM-score ≥ 0.3, ≥30% sequence identity, bidirectional coverage), and every pool entry that shared a cluster with any validation or test entry was removed. Second, using a single exhaustive all-against-all MMseqs2 search over the same combined set and the homology criteria described above (≥30% sequence identity, ≥80% coverage of either sequence, E ≤ 10⁻³), we removed every pool entry that was homologous to any validation or test entry. Pool entries whose structures could not be indexed by Foldseek were also removed, as they could not be checked for leakage. The final whole-BMRB training set contained 12,054 entries.

### SLAE-CS and SLAE-ERR Architecture

SLAE-CS was developed based on SLAE protein structure encoder and finetuned for chemical shift prediction task^6^. The input structure from AlphaFold2, was first converted into a graph representation (8 Å radius cutoff, atom37 featurization) and passed through the pre-trained SLAE to obtain a per-residue embedding of dimension 128. A prediction trunk was trained on top of these SLAE embeddings, which started with a linear projection (128 ⟶ 256), followed by a depth-wise 1D convolution (kernel size 5) to incorporate local sequence context. The resulting features were then processed by two TransformerEncoderLayers (four attention heads, feed forward dimension 1024) to capture longer range dependencies, followed by four residual feed forward blocks, each consisting of LayerNorm, a GEGLU expansion (256 → 512), and a linear projection with a residual connection. A final linear layer (256 → 1) outputed the predicted chemical shift μ, with its bias initialized to the empirical prior mean (^15^N = 119.0 ppm). The model was trained by minimizing the Huber loss between the predicted μ and the ground truth chemical shift, using RMSE as the selection metric (Fig. S3).

Discriminative learning rates were used: the prediction trunk was optimized with a learning rate of 3×10^-5^, while the encoder was finetuned at 3×10^-4^. Both parameter groups were optimized using AdamW with a weight decay of 1×10^-4^. The learning rate followed a linear warmup cosine annealing schedule, with warmup steps set to min(100, total training steps/10) and a warmup starting learning rate of 1×10^-6^. The gradients are clipped to a maximum norm of 1.0. All models were trained in FP32 precision with seeds 1–10 for the ten ensemble members, and cuDNN determinism was enforced (benchmark mode disabled, deterministic algorithms required) to ensure reproducibility across runs (Fig. S3).

SLAE-ERR shared the SLAE-CS’s architecture but was trained and applied independently, taking as input the SLAE’s per-residue embeddings. The error trunk was trained to predict the absolute residual, |error|, rather than the signed residual; this choice was motivated in part by the approximately symmetric, unbiased distribution of signed per-residue error observed empirically (median = −0.005 ppm; Fig. S5c), which suggested few systematic, direction-dependent patterns for a model to learn, making the unsigned error magnitude a more tractable and informative target. To incorporate ensemble uncertainty, this embedding was augmented with features from a 10-member deep ensemble of independently seeded prediction-trunk outputs: the per-residue standard deviation across the ensemble (one channel), and the 10 per-residue predictions themselves, sorted in ascending order and mean-centered (10 channels). Sorting renders the input invariant to the arbitrary ordering of the exchangeable ensemble members; centering removes the ppm-scale offset so these features are numerically comparable to the embedding. Because the target (the absolute residual) is non-negative, the final layer’s bias was initialized to the mean absolute out-of-fold residual of the training set, computed at run time. The trunk was trained with Huber loss (δ = 1.0) jointly with encoder fine-tuning (lr = 3×10^-4^ encoder, 5×10^-4^ error trunk; AdamW, weight decay 1×10^-4^; linear warmup/cosine decay; 10 epochs; seed=42), selecting the checkpoint with the highest validation Spearman correlation between predicted and observed |error| (Fig. S3).

Because SLAE-ERR was trained on prediction errors estimated from the training set, we computed these training-set residuals out-of-fold to prevent data leakage. Specifically, the training set was partitioned into five folds while keeping sequence/structure similarity clusters intact, with clusters assigned using size-descending bin packing. For each fold, residuals were predicted using ensemble members retrained on the remaining four folds. Validation and test residuals were computed directly using the full ensemble. For both models, batch size 1 was used during inference for evaluation.

### Train size sweep setup

To assess how SLAE-CS performance scales with training-set size, we retrained the model on subsets of the training data for each of the four referencing schemes (PANAV, LACS, NOREF, and NOREF_whole). For each scheme, we randomly shuffled the training-set identifiers once using a fixed random seed. We then selected the first 10%, 20%, …, 90% of the shuffled list to create training subsets of increasing size. This nested sampling strategy ensures that each smaller subset is contained within every larger subset, so that training-set size, rather than data composition, is the primary variable across the learning curves. Identical nested subsets (465–4,182 entries) were used for all PANAV, LACS and BASELINE schemes. Whole BMRB uses a larger training set (12,054 entries), from which nested subsets of 1,205–10,849 entries were generated in the same manner.

Each subset was used to train a single model with random seed 1, keeping the architecture, optimizer, learning-rate schedule, number of epochs, loss function and all others identical to those of the corresponding full-data models (see SLAE-CS Architecture). The validation and test sets were fixed across all subset sizes and referencing schemes, using the PANAV validation set (385 proteins) and test set (413 proteins). All models were evaluated with inference performed at batch size 1. The 100% point on each learning curve corresponds to the full-data model trained with the same random seed (seed 1).

### Other foundation models setup

#### ESM3

We used ESM3-open (esm3-sm-open-v1; 48 layers, d = 1536). Each chain was given to the model as its sequence plus the AF2 backbone coordinates. No secondary-structure, SASA or function annotations were supplied, and per-residue confidence conditioning was set to 1. The per-residue embedding is the final-layer hidden state (1536-dimension) with the BOS/EOS positions removed. In the sequence-only variant, coordinates were omitted.

#### MACE-OFF23

Per-residue descriptors were computed with MACE-OFF23 (large; r_max = 5 Å, two interaction layers), following Bojan et al.^5^. For each residue, a local environment was cropped from the protonated AF2 model: every residue with any atom within 5 Å of the focus residue. MACE was evaluated on this all-atom environment. The invariant (scalar) node features of both interaction layers at the backbone amide nitrogen were concatenated, giving a 448-dimension embedding.

#### ProteinMPNN

We used the encoder of vanilla ProteinMPNN (v_48_020; 3 message-passing layers, 48 nearest neighbors, hidden size 128) and discarded the sequence decoder. Its input is the backbone N/Cα/C/O coordinates plus an ideal virtual Cβ, together with residue indices for the relative positional encoding; the amino-acid sequence is not used. The 128-dimension node embeddings were used in fine-tuned settings: the same encoder, initialized from the pretrained weights, was applied on the fly to each batch and trained end-to-end with prediction trunk.

#### Training

For the frozen embeddings (ESM3, ESM3 sequence-only, MACE-OFF23), each embedding dimension was transformed via z-score using the mean and standard deviation over all training residues. These statistics were stored with the model and applied unchanged to validation and test. Only prediction trunk was trained, and its learning rate was selected as: 3e-4 for both ESM3 variants and 3e-5 for MACE-OFF23. Fine-tuned ProteinMPNN was trained like SLAE-CS, with separate learning rates for the encoder (1e-3) and the trunk (3e-5). All models were trained with 10 random seeds and evaluated as for SLAE-CS.

### Protein structure preparation

Protein structure models in AlphaFold2 were generated using ColabFold^72^: colabfold_batch --amber --templates --num-recycle 3 --use-gpu-relax --jobname-prefix <job_name> --zip <fasta> <output>

Structures were predicted with template usage enabled and three recycling iterations, followed by Amber relaxation on GPU. Relaxed structures were saved for each entry.

### Tag and linker masking

Many BMRB-deposited sequences include non-native residues introduced during recombinant expression and purification, such as polyhistidine affinity tags, protease cleavage sites (e.g., TEV, thrombin), and short glycine/serine linkers, which do not reflect genuine biological sequence context and can introduce spurious signal if left unmasked. We applied a sequence-based heuristic labeling scheme to flag such regions: each residue in a sequence was assigned one of four labels: A, a standard amino acid position; P, a proline, excluded separately from ^15^N predictions and error calculations since proline lacks a backbone amide hydrogen; T, a residue within a recognized affinity-tag or cleavage-site motif; and t, a residue in the flanking region immediately adjacent to such a motif. This labeling was applied independently to every entry in the curated dataset (6,123 sequences for rBMRB and 12,054 sequences for additional whole BMRB training set), based solely on primary sequence, with no dependence on experimental chemical-shift assignments. The resulting per-residue tag/flank labels were used as an mask during model training and evaluation, allowing tag- and linker-derived residues to be excluded where appropriate.

### Model ensembling

Before comparing the three re-referencing conditions, we found that random seed choice substantially influenced chemical shift prediction performance. To reduce this variance and enable a more reliable analysis of the models’ understanding of local chemical environments, we employed deep ensembles^73^, training ten models in parallel for each condition and averaging their predictions for evaluation; this stabilized both RMSE and Kendall’s τ as more seeds were averaged, and introduced large improvement at the same time (Fig. S4a). By comparison, using Monte Carlo dropout method^74^ did not achieve the same effect (Fig. S4b). All evaluations are based on ensemble predictions unless otherwise noted.

### Protein-level bootstrap

Residues from the same protein are not statistically independent: they share chemical-shift referencing, sample conditions, structure quality and assignment practice, so their prediction errors are correlated. Treating residues as independent observations would thus inflate significance. We estimated uncertainty with a cluster bootstrap that uses the protein, rather than the residue, as the resampling unit. In each resample, N proteins were drawn with replacement from the N proteins in the analysis set, all residues of each drawn protein were pooled, and the statistic of interest (a difference in median |error| or an AUROC) was recomputed. For AUROC, the out-of-fold classifier scores were held fixed and only the evaluation set was resampled. 95% confidence intervals are the 2.5th and 97.5th percentiles of the bootstrap distribution. Two-sided P values are twice the smaller of the fractions of resampled statistics ≤ 0 and ≥ 0; when no resample crossed zero, P is reported as P < 1/B (B is resample times).

### Filtering of pairs for intrinsic noise analysis

To estimate how much of the disagreement between independent chemical shift depositions of the same protein is attributable to a fixable referencing offset rather than genuine measurement scatter, we identified all BMRB entries sharing an identical amino acid sequence and formed all pairwise comparisons within each such cluster (criteria of ΔpH < 0.2 and Δtemperature < 2.5K are applied here, see Figure S6).

Pairs were then screened using a three-stage procedure. First, an automated classifier parsed each entry’s assembly name, entry title, and assigned-shift-list name tags where present in NMR-STAR file, together with the structured “Physical_state” and “Thiol_state” fields. This screening used keywords indicating changes in binding partners, oligomeric states, redox or thiol states, or physical states (denatured versus native); any pair flagged by these criteria was excluded.

Second, because BMRB metadata are inconsistently formatted and frequently omits relevant information, every pair the classifier left unflagged was manually inspected. Pairs confirmed to differ in binding state, oligomeric state, redox or thiol state, post-translational modification (e.g., phosphorylation, glycosylation, methylation), covalent modification, point mutation, or documented ligand were excluded from the intrinsic-noise estimate (n = 95 pairs). Candidate pairs whose documented difference was a minor buffer, salt, sample-concentration variation, existence of multiple conformers under same condition, or for which no state difference could be identified in either metadata or the original publication, were retained. Lastly, pairs whose re-referenced chemical shifts agreed to within 0.01 ppm RMSE were treated as near-duplicate depositions and excluded. The complete pair-by-pair disposition is provided in Supplementary Material Table. S2 and S3.

### Decomposition of inter-deposition disagreement into referencing offset and intrinsic noise

For each condition-consistent pair and each re-referencing method (LACS, PANAV), we computed the per-residue chemical shift difference between the two depositions and decomposed the resulting root mean squared disagreement into a referencing offset term and an intrinsic, reference-independent noise term. Offset was defined as the difference between the two depositions’ method-specific offsets (offsetA – offset; from PANAV or LACS); and intrinsic RMSE as the root mean squared difference after subtracting this offset from the raw per-residue differences. Aggregate offset and intrinsic RMSE for each method were computed as the pooled root-mean-square across all condition-consistent pairs.

### Statistical testing of secondary-structure and burial effects on prediction error

Effects of secondary structure and solvent accessibility on ^15^N prediction |error| were assessed using a protein-cluster bootstrap. For a given contrast between two groups, we computed the observed difference in medians and generated its null distribution by resampling proteins with replacement 20,000 times (bootstrap replicates yielding fewer than 5 residues in either group were discarded).

To isolate secondary-structure effects from burial effects, we restricted comparisons to same-factor contrasts within the nine secondary structure-solvent accessibility groups, comparing secondary structure classes with fixed solvent accessibility, and solvent accessibility within a fixed secondary structure class, excluding contrasts that varied both factors simultaneously, and applied Holm–Bonferroni correction across the resulting family of nine tests. Figure 5a displays the three same-secondary-structure, cross-solvent-accessibility contrasts from this family (buried versus exposed, tested separately within helix, sheet and loop residues).

For the systematic analysis of burial effects (Fig. 5e), we compared median |error| between buried and exposed residues across 79 strata: the full test set, individual secondary structure classes, individual residue types, and residue type x secondary structure combinations. Only strata with at least 30 residues in each burial class were retained. For each stratum, significance was assessed using two tests: a protein-cluster bootstrap (100,000 resamples) and a two-sided Mann–Whitney U test on residue-level |error| values. P values from each test were Bonferroni-corrected across all 79 strata (P_adj_ = min(P × 79, 1); per-test significance threshold, P < 1.27 × 10^-4^). This correction controls the family-wise error rate rather than the false discovery rate. A stratum was considered significant only if both tests yielded P_adj_ < 0.01 and the 95% bootstrap confidence interval excluded zero.

To assess whether the burial effects identified for SLAE-CS were specific to this model, we applied the same analysis pipeline—including the 79 strata, residue-count filter, statistical tests, and bootstrap settings—to the 10-seed ensemble-mean predictions of ESM3 (structure + sequence), ESM3 (sequence only), MACE-OFF23, and ProteinMPNN (Fig. S8). Because the comparisons focused on the K = 10 strata identified as significant in the SLAE-CS analysis (Fig. 5e), Bonferroni correction was applied across these ten comparisons for each model (P_adj_ = min(P × 10, 1); per-test significance threshold, P < 1 × 10^-3^). The same significance criteria were applied.

### Selection of proteins for molecular dynamics simulation

Molecular dynamics trajectories were generated for 48 proteins drawn from the SLAE-CS test set. These proteins span 80–254 residues (median 118 aa), each with near-complete experimental backbone-^15^N chemical-shift coverage (63–236 observed, non-proline residues per protein, corresponding to 59–98% of the full chain, median 90%). All proteins have a mixed secondary-structure composition, containing both β-sheet (9–54% of observed residues, median 30%) and α-helix (13–55%, median 32%) content, providing within-protein structural contrast for the downstream analysis. Starting structures were taken from AlphaFold2 models with high per-protein confidence (median pLDDT 85.8–98.5). Experimental conditions were near-neutral (pH 4.5–8.0, median 6.8). Coverage, secondary-structure, and confidence metrics were computed directly from the experimental/AlphaFold2 record for each protein and do not depend on the chemical-shift predictor’s output.

### Molecular dynamics simulations setup

All-atom explicit-solvent MD simulations were performed with GROMACS 2022.2^75^ using the AMBER99SB-ILDN force field^76^ and the TIP3P water model^77^. Non-protein heteroatoms were removed from the starting structures and hydrogens were added with pdb2gmx. Each protein was solvated in a rhombic dodecahedral box with a minimum solute–edge distance of 1.0 nm, neutralized and brought to 0.1 M NaCl. Systems were energy-minimized by steepest descent (F_max_ < 1000 kJ mol^-1^ nm^-1^), then equilibrated with heavy-atom position restraints for 100 ps in the NVT ensemble and 500 ps in the NPT ensemble. Production simulations of 1 µs were run in the NPT ensemble at 298 K (velocity-rescaling thermostat, τ_T_ = 1.0 ps) and 1 bar (C-rescale barostat, τ_P_ = 5.0 ps) with a 2-fs time step and LINCS constraints on bonds to hydrogen. Short-range van der Waals and electrostatic interactions were truncated at 1.0 nm, long-range electrostatics were treated with PME, and long-range dispersion corrections were applied to energy and pressure. The first 100 ns of each trajectory were discarded, and the remaining 901 ns were sampled every 1 ns (901 frames per protein) for analysis.

### Backbone order parameter (S^2^) from MD trajectories

For each protein, the backbone amide N–H order parameter was computed using the isotropic reorientational eigenmode dynamics (iRED) formalism^78^, which is alignment-free: rather than requiring superposition onto a reference structure, iRED derives internal order parameters directly from pairwise dot products of instantaneous N–H bond vectors within each trajectory frame, avoiding artifacts introduced by the choice of alignment reference. For each frame, unit N–H bond vectors e_i_ were computed for all qualifying backbone amides, and a reorientational covariance matrix was constructed as

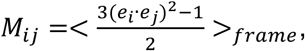

averaged over all frames. This matrix was diagonalized, and the five eigenmodes with the largest eigenvalues, corresponding to overall isotropic tumbling, were used to compute each residue’s order parameter as

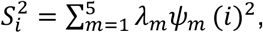

where *λ_m_* and *ψ_m_* are the eigenvalue and eigenvector of mode m. This calculation was performed once per protein using every qualifying backbone amide simultaneously. Residues lacking an amide hydrogen were excluded.

### Backbone dihedral (ψ) circular variance from MD trajectories

Backbone ψ dihedral angles (N_i_–C^α^_i_–C_i_–N_i+1_) were computed for every frame using MDTraj^79^. For each residue, the circular variance of ψ across the trajectory was defined as

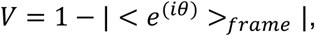

where the average is taken over all sampled dihedral angles *θ*; V ranges from 0 (rigid, non-fluctuating dihedral) to 1 (no preferred angle). ψ depends only on its four defining atoms, while S^2^ is calculated from dot products within each frame. Neither requires structural alignment.

### Residue selection for MD-based loop analyses

The loop analyses in Fig. 5c, d and f used residues which had experimental and predicted backbone ^15^N shifts available (Pro excluded) and which retained the same secondary structure and solvent accessibility status for most of the MD simulation. Static labels were taken from the AF2-predicted structure. Secondary structure and solvent accessibility were assigned based on the same rationale as below “Extracting structural features”. For each of the 48 trajectories, the first 100 frames (100 ns) were discarded as equilibration and the remaining 901 frames (1-ns spacing) were analyzed with MDTraj^79^. In every frame, the DSSP class and amide N solvent accessibility were recomputed with the same definitions. For each residue, we calculated the fraction of frames in which its DSSP class matched the static loop label and the fraction in which its burial state matched the static solvent accessibility. A residue was kept when both fractions were ≥0.5. This 50% threshold retained 351 buried and 900 exposed loop residues. Fig. 5c additionally required ψ(i) to be defined, and the per-protein correlation included proteins with at least five such residues in a burial class. Fig. 5d and f additionally required both φ(i) and ψ(i) to be defined.

### Ramachandran region assignment

Backbone φ(i) and ψ(i) dihedrals of the selected residues were computed with MDTraj for every post-equilibration frame. Ramachandran regions followed Hollingsworth and Karplus^45^. We digitized the boundaries of eight regions (β, P_II, ζ, γ′, δ, α, δ′ and P_II′) from their Fig. 6A with WebPlotDigitizer^80^ and fitted an ellipse to each boundary by least squares (scikit-image^81^). Fitting and assignment were performed in the shifted coordinate frame of that figure (φ from 0° to 360°, ψ from −120° to 240°), in which each region forms a single closed contour. Each frame was assigned to the region whose ellipse contained it. The α region, which is nested within δ, took precedence over δ; other overlaps were assigned to the ellipse in which the point had the smallest normalized radius. Frames inside no ellipse were assigned to the region with the nearest ellipse boundary if that distance was ≤30°, measured as the Euclidean distance in degrees with periodic images; the remaining 3.0% of frames were left unassigned. The occupancy of a residue in each region was the fraction of its frames assigned to that region.

### Error projection onto the Ramachandran plane (Fig. 5d)

Each residue carries a single absolute error, which was distributed over the (φ, ψ) positions of its frames. Every frame was weighted by 1/901, so each residue contributed equal total weight. For each burial class, frame weights and error-multiplied frame weights were accumulated in 2° × 2° bins and smoothed with a periodic Gaussian kernel (σ = 6°). The color shows the ratio of the two smoothed maps, which is the time-weighted mean error of the residues that visit each (φ, ψ) position. Because each residue has one experimental shift, the map shows where poorly predicted residues spend their time, not a per-frame error. The color scale, shared by both panels, spans the 2nd–98th percentiles of the values inside it.

### Extracting structural features

Secondary structure and solvent accessibility, the two basic structural descriptors underlying the analyses in Fig. 5, were computed directly from each AlphaFold2-predicted test set structure. Secondary structure was assigned using DSSP^82^ (via mkdssp^83^), with the eight-class DSSP code collapsed into three categories: helix (H, G, I), sheet (E, B), and coil (all other codes). Solvent accessibility of each backbone amide was classified using a structure-based, DSSP-independent criterion: the solvent-accessible surface area (SASA) of the backbone amide nitrogen and its attached hydrogen was computed using the Shrake–Rupley algorithm^84^ in Biopython^85^ (1.4 Å probe radius, 960 points per atom), and residues with combined N–H SASA greater than 0 Å² were classified as exposed and all others as buried; prolines were excluded from this classification.

We also computed a broader set of structural features from AF2 structures and assessed their influence on where ^15^N prediction fails. We calculated more than 20 features spanning the principal known determinants of backbone amide ^15^N chemical shifts, including: backbone dihedral angles at the target residue (φ, ψ) and at its immediate sequence neighbors (ψ of residue i−1, φ of residue i+1), the side-chain χ_1_ rotamer, neighboring-residue identity^86^, a preceding-proline flag, the amide donor hydrogen bond directly involving the predicted N–H (distance, donor/acceptor angles, and a Kabsch–Sander-style electrostatic energy term, computed using the geometric criteria of SPARTA+^21^), the ring-current contribution at the amide nitrogen from a Haigh–Mallion ring current model^22,87^, and local heavy-atom packing density within 8 Å of the nitrogen.

Beyond these explicit descriptors, we incorporated discrete structural tokens, the Foldseek 3Di alphabet^69^ and the neighborhood-aware 3Dn alphabet^88^, which compress nearest neighbor and complex local geometry into a small vocabulary, reducing dimensionality and helping expose the patterns underlying prediction accuracy. We also included two dynamics indicators, p(exchange) from Dyna-1 and the order parameter S^2^ from TALOS-N (through makeshift), given their established correlation with chemical shifts^15,89^. pLDDT from AF2 output^53^ and the experimental conditions pH and temperature^90,91^ were also collected.

### Grouping structural features for permutation importance

Because several features are highly correlated or are alternate representations of the same underlying quantity (e.g., the sine and cosine components of a single dihedral angle), naively permuting individual features can underestimate their true importance. We therefore grouped features prior to permutation testing, using a two-step, largely data-driven procedure.

First, we constructed a unified pairwise association matrix encompassing both categorical and numerical features, with all association values scaled to the range [0, 1]: for categorical–categorical pairs, bias-corrected Cramér’s V^92^ (appropriate for the high-cardinality categorical features, such as residue identity and the Foldseek 3Di/3Dn tokens); for numeric–numeric pairs, the absolute Spearman correlation; and for categorical–numeric pairs, the correlation ratio η, which quantifies the fraction of numeric variance explained by categorical group membership. Features were then hierarchically clustered (average linkage, distance = 1 − association), and empirical groups were defined by cutting the resulting dendrogram at an association threshold of 0.7 (Fig. S15).

This purely correlation-based clustering cannot recognize that a dihedral angle’s sine and cosine components jointly encode a single, non-monotonic (circular) quantity. We therefore additionally enforced a small set of “must-link” constraints via union-find, requiring the sine and cosine of each dihedral (φ, ψ, ψ of residue i−1, φ of residue i+1, χ_1_), and all five amide hydrogen-bond descriptors (distance, donor and acceptor angle cosines, existence, and energy) to remain in the same group. Combining these must-link constraints with the empirical clusters above yielded the final feature groups used for grouped permutation importance (Fig. 6c): backbone conformation (3Di token, secondary structure class, and the ψ dihedrals of residue i and i−1), local packing density (3Dn token and pack_N8), hydrogen bond geometry (the five amide hydrogen-bond descriptors), the φ dihedral, the χ_1_ dihedral, the φ dihedral of residue i+1, previous- and next-residue identity, and the remaining features retained as individual groups: residue identity, solvent accessibility, the order parameter S^2^, p(exchange), pH, temperature, model confidence (pLDDT), and the ring current effect at the amide nitrogen.

### Grouped permutation importance for hard-versus-easy classification

To identify which feature groups (defined above) carry the most information about where prediction fails, we quantified their contribution to the hard-versus-easy classification task via grouped permutation importance. Residues were labeled “easy” if their |error| fell at or below the 50th percentile and “hard” if it fell at or above the 90th percentile across the full test set; residues in between were excluded from this analysis. A gradient-boosted classifier (HistGradientBoostingClassifier, scikit-learn^93^; maximum depth 3, learning rate 0.05, 400 boosting iterations, L2 regularization 1.0, balanced class weighting) was trained to discriminate hard from easy residues using categorical features (one-hot encoded) and standardized numeric features as input. Model performance was evaluated using k-fold cross-validation (5-fold), with folds defined at the protein level so that no protein’s residues appeared in both the training and held-out test fold within a given split.

For each fold, we first computed the baseline out-of-fold AUROC using the trained classifier’s predictions on the held-out test residues. For each feature group, we then jointly permuted the values of all columns belonging to that group across the held-out residues (preserving the joint structure within a group, e.g., a dihedral’s sine and cosine components were permuted together using the same permutation index), recomputed AUROC on the permuted data, and recorded the drop relative to the baseline; this was repeated six times per group per fold using independent random permutations, and averaged. The final importance of each feature group was defined as the mean drop in AUROC across all five folds, with the standard deviation across folds reported as an uncertainty estimate (Fig. 6c).

### Outlier identification

We identified proteins whose error SLAE-ERROR misjudged. We fit a linear regression of mean ground-truth |error| on mean predicted |error| across all proteins. For each protein, the residual is its mean ground-truth |error| minus the fitted value. We then flagged outlying residuals using the interquartile range (IQR). Let Q1 and Q3 be the 25th and 75th percentiles, and IQR = Q3 − Q1. Proteins with a residual above Q3 + 1.5 × IQR were labelled under-predicted outliers: their actual error is larger than the model expects. Proteins with a residual below Q1 − 1.5 × IQR were labelled over-predicted outliers. This rule gave 18 under-predicted and 0 over-predicted outliers (Fig. 6e).

### Oracle lower bound of the SLAE-ERR error

To place the MAE of SLAE-ERR in context, we estimated the lowest MAE that a predictor of the absolute error could reach given experimental noise (an “oracle” error predictor). We wrote the observed signed error of residue *i* as

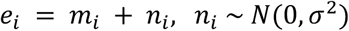

where *m_i_* is the true error of the model and *n_i_* is independent Gaussian experimental noise. We set *σ* = 0.5 ppm, based on the median per-pair intrinsic RMSE of 0.48 ppm (Fig. 3c). The oracle knows |*m_i_*| and *σ* but not *n_i_*, so it cannot predict the observed |*e_i_*| exactly. Under absolute loss its best prediction is the median of |*m_i_* + *n_i_*|, and its expected absolute error is

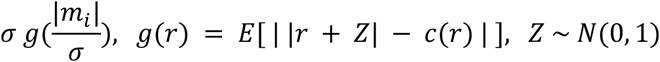

where *c*(*r*), the median of |*r* + *Z*|, solves *Φ*(*c* − *r*) − *Φ*(−*c* − *r*) = ^1^, and *Φ* is the standard normal cumulative distribution function. We evaluated *g* numerically. Because *m_i_* is not observed, we estimated |*m_i_*| as |*e_i_*|. The bound is the mean over the residues used to score SLAE-ERR,

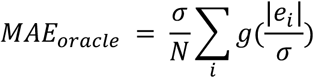

with *N* = 42153 residues from 407 test-set proteins. This gave 0.348 ppm (95% CI 0.347–0.349) compared with an MAE of 0.760 ppm (95% CI 0.739–0.780) for SLAE-ERR. Confidence intervals are from a protein-level bootstrap with 2,000 resamples.

