## Supplementary Material for "Learning the limits of protein structure models with NMR chemical shifts"

Cizhang Zhao[1], Gina El Nesr[2], Yilin Chen[3], and Hannah K. Wayment-Steele[1]

[1]Department of Biochemistry, University of Wisconsin-Madison, Madison, WI, USA

[2]Department of Integrated Structural and Computational Biology, The Scripps Research Institute & Howard Hughes Medical Institute, La Jolla, CA, USA

[3]Department of Bioengineering, Stanford University, Stanford, CA, USA

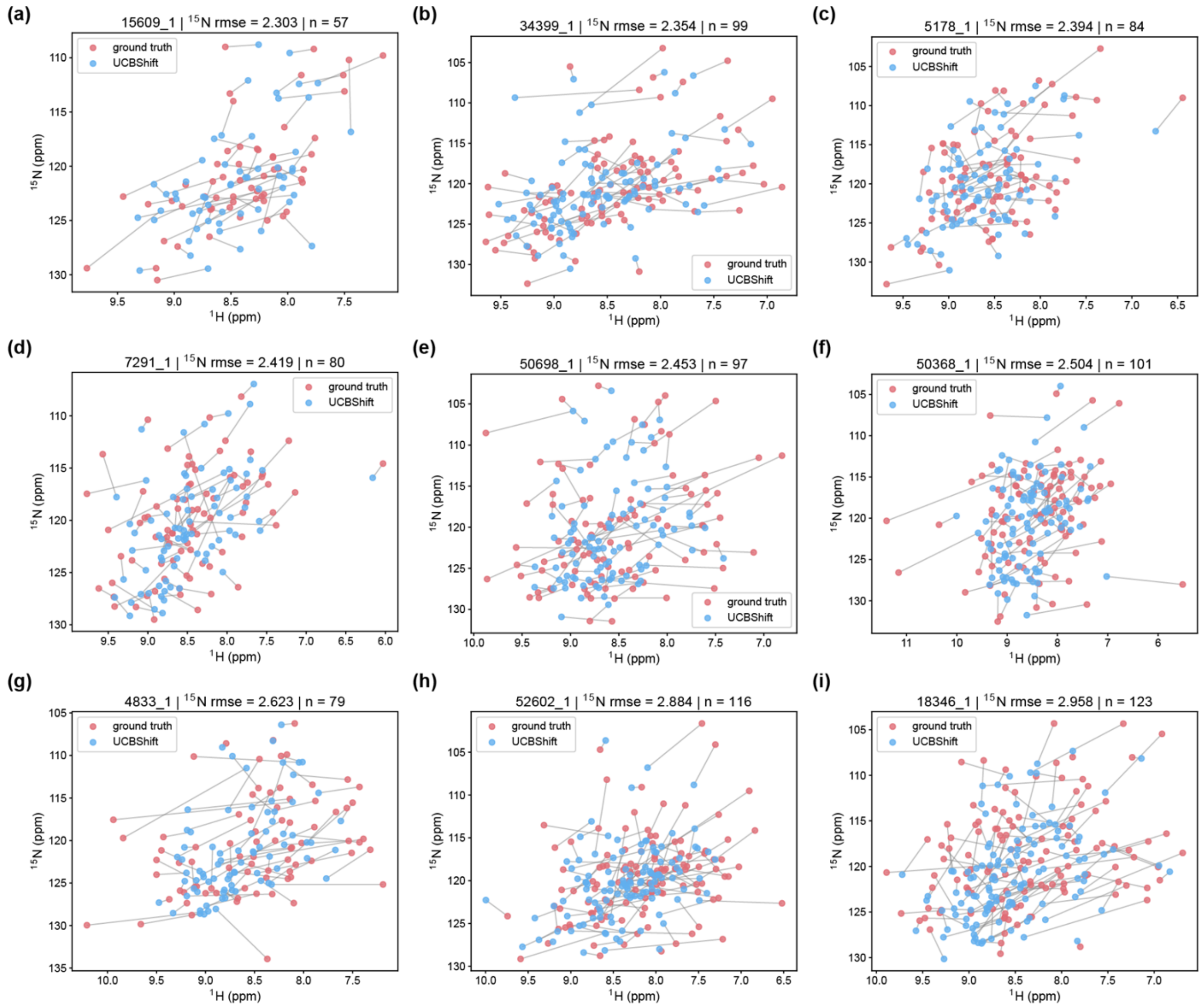


**Fig. S1 | Backbone $^{15}$N chemical shifts are poorly predicted.**

a–i, Simulated $^{1}$H–$^{15}$N HSQC spectra comparing experimental (red) and and predictions from near state-of-the-art UCBshift (blue) backbone amide chemical shifts for nine randomly selected proteins. Lines connect the experimental and predicted peaks of the same residue. Each panel title gives the BMRB entry ID and the $^{15}$N RMSE (ppm) and the number of residues with both $^{1}$H and $^{15}$N shifts (n). Panels are ordered by increasing $^{15}$N RMSE (2.30–2.96 ppm).

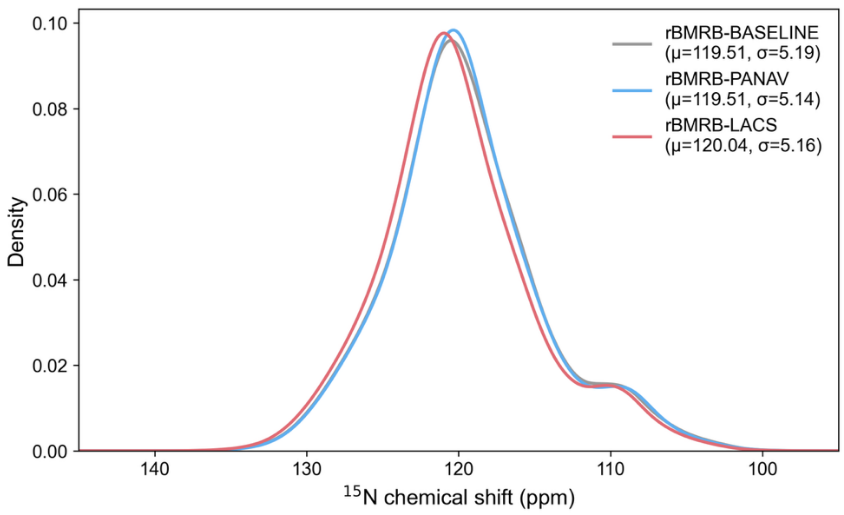


**Fig. S2 | Distribution of backbone $^{15}$N chemical shifts under different re-referencing schemes.**

Kernel density estimates of backbone amide $^{15}$N chemical shifts under each re-referencing condition. The distributions are nearly superimposable: rBMRB-BASELINE ($\mu$ = 119.51 ppm, $\sigma$ = 5.19 ppm), rBMRB-PANAV ($\mu$ = 119.51 ppm, $\sigma$ = 5.14 ppm) and rBMRB-LACS ($\mu$ = 120.04 ppm, $\sigma$ = 5.16 ppm).

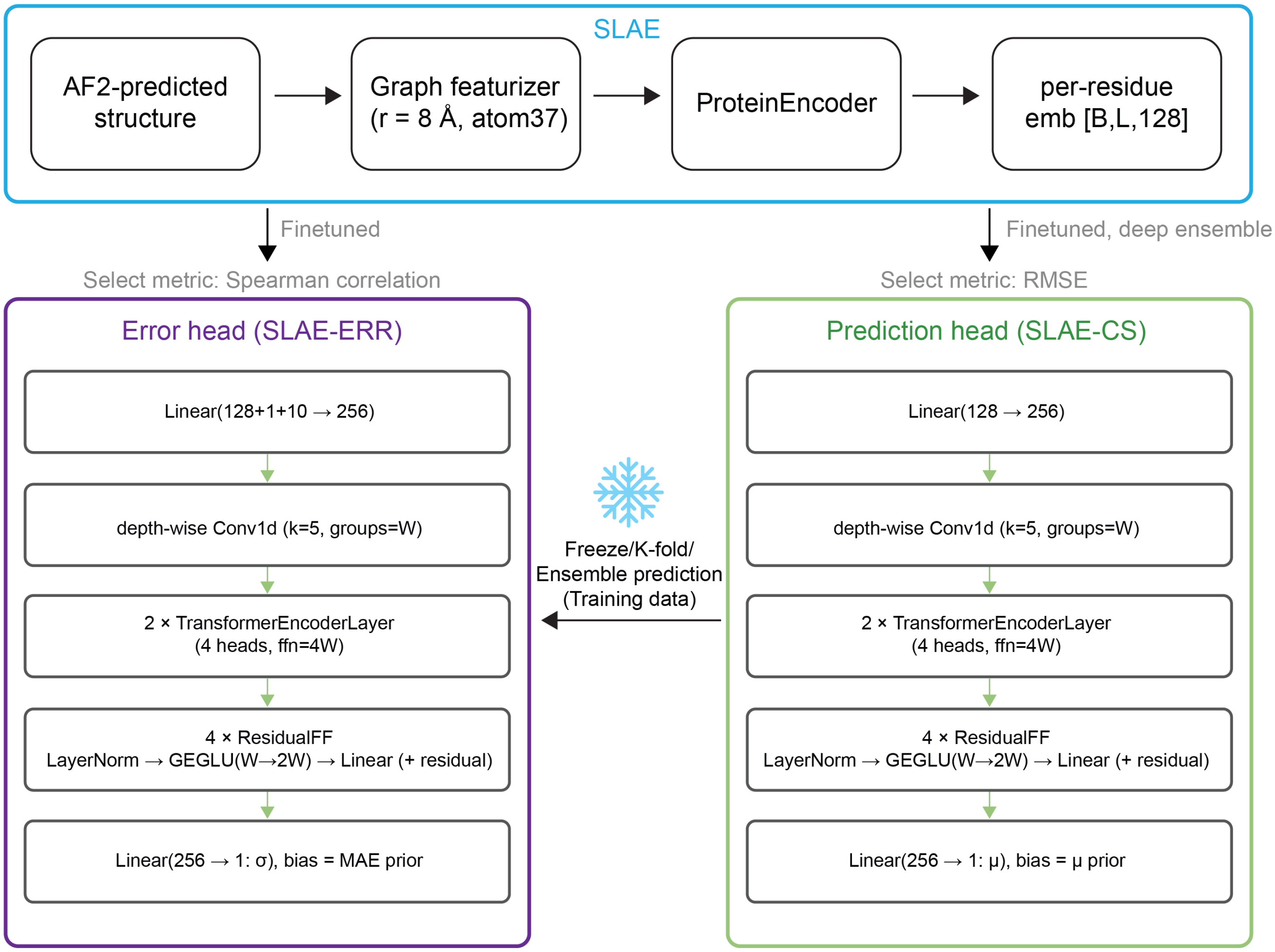


**Fig S3 | Architecture of SLAE-CS and SLAE-ERR.**

Structures are encoded using SLAE and output embeddings are further used for downstream predictions. SLAE parameters are finetuned in the training process. Both heads for chemical shift and error predictions share very similar architecture except the initialized bias, but error prediction uses the outputs of frozen prediction heads as training data (snowflake indicates frozen model). Prediction of training set is acquired through K-fold training (5 folds in total). The training of two heads does not share any parameters.

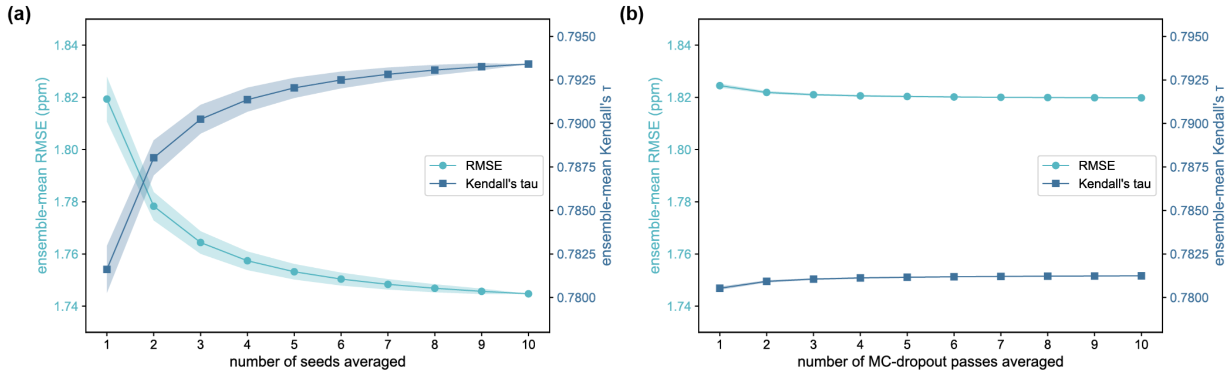

**Fig. S4 | Ensembling over independently trained seeds, but not Monte Carlo dropout passes, reduces prediction error and improves ranking performance.**

Results are shown for SLAE-CS-PANAV. (a) ensemble mean RMSE (left axis, cyan) decreases and ensemble mean Kendall's τ (right axis, navy) increases as more independently trained model seeds are averaged, plateauing beyond approximately six seeds. All subsequent analyses use ten seed ensembles. (b) By contrast, averaging over an increasing number of Monte Carlo dropout passes within a single trained model yields only marginal improvements in both RMSE and Kendall's τ, indicating that seed-to-seed variance, rather than dropout-induced stochasticity, is the dominant source of epistemic uncertainty captured by this ensembling strategy. Shaded band, s.d. across ensembles of the same size.

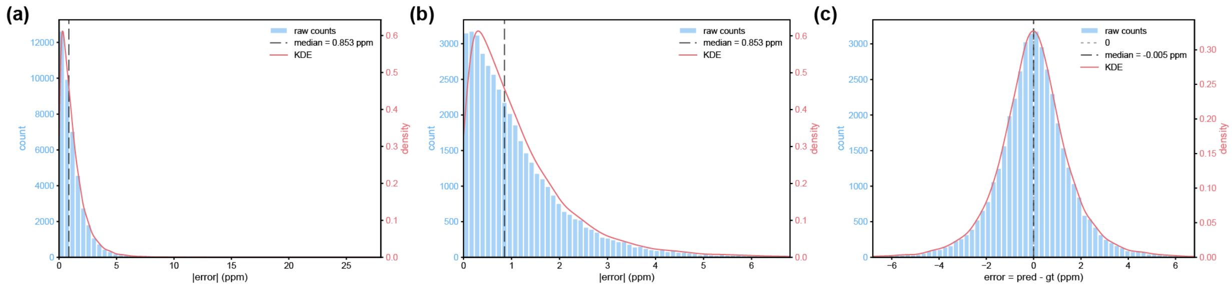


**Fig. S5 | Per-residue $^{15}N$ prediction errors are unbiased but heavy-tailed, and rare large outliers inflate RMSE.**

(a) Distribution of per-residue |error| across the full PANAV test set (n = 42,153 residues; 10-seed ensemble predictions), shown over the complete data range. The distribution is strongly right-skewed: most residues are predicted with small error (median = 0.853 ppm), while a long tail of infrequent but very large errors extends out to ~28 ppm. (b) The same distribution, restricted to the 99.5th percentile (|error| < ~6.8 ppm) to better resolve the shape of the bulk of the distribution; median (dashed line) is unchanged. (c) Distribution of signed per-residue error (predicted minus ground truth, same restriction as b), which is approximately symmetric around zero (median = −0.005 ppm), indicating that the model is neither systematically over- nor under-predicting chemical shifts. In all panels, blue bars show raw counts (left axis) and the red curve shows a Gaussian kernel density estimate (right axis).

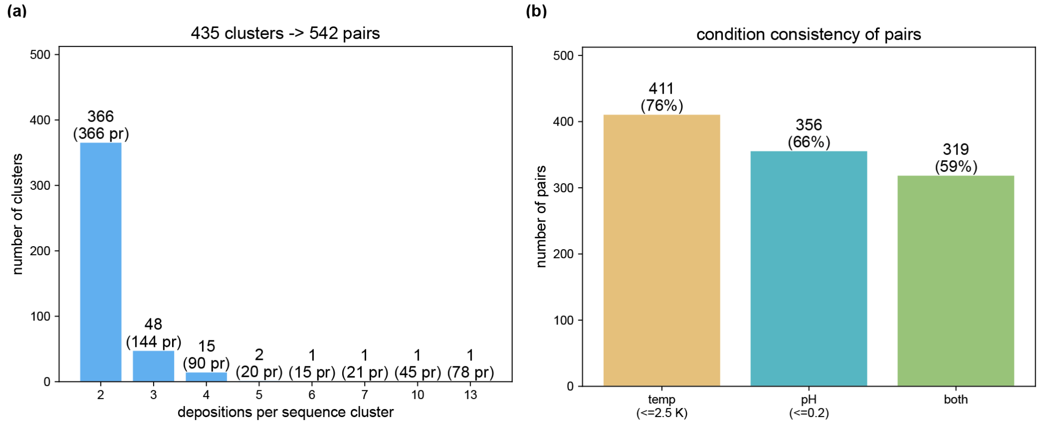


**Fig. S6 | Construction of sequence-matched deposition pairs used to estimate inherent measurement noise.**

(a) Distribution of cluster sizes among rBMRB protein entries sharing an identical sequence (depositions per cluster), with the number of pairwise comparisons each cluster size contributes shown in parentheses. Clustering the 7,758-entry rBMRB (BASELINE dataset, before finding common entries with PANAV and LACS) by sequence identity identified 435 clusters containing two or more independent depositions, yielding 542 candidate pairs with at least five co-observed $^{15}N$ chemical shifts.

(b) Number of these pairs retained after requiring that paired depositions share consistent experimental temperature (≤2.5 K), pH (≤0.2), or both, isolating pairs whose residual disagreement reflects intrinsic measurement noise rather than condition-driven chemical shift variation.

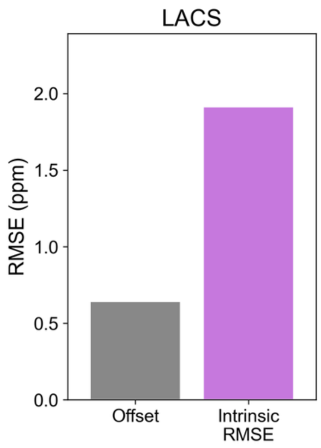


**Fig.S7 | Intrinsic RMSE exceeds the referencing offset about three-fold under LACS re-referencing, as under PANAV**

Aggregate referencing offset and intrinsic RMSE (pooled root mean square) by referencing method LACS (n = 51). The intrinsic noise (1.91 ppm) is roughly three-fold larger than the corresponding referencing offset (0.64 ppm), indicating that most of the disagreement between sequence-identical depositions likely reflects irreducible noise for predictors.

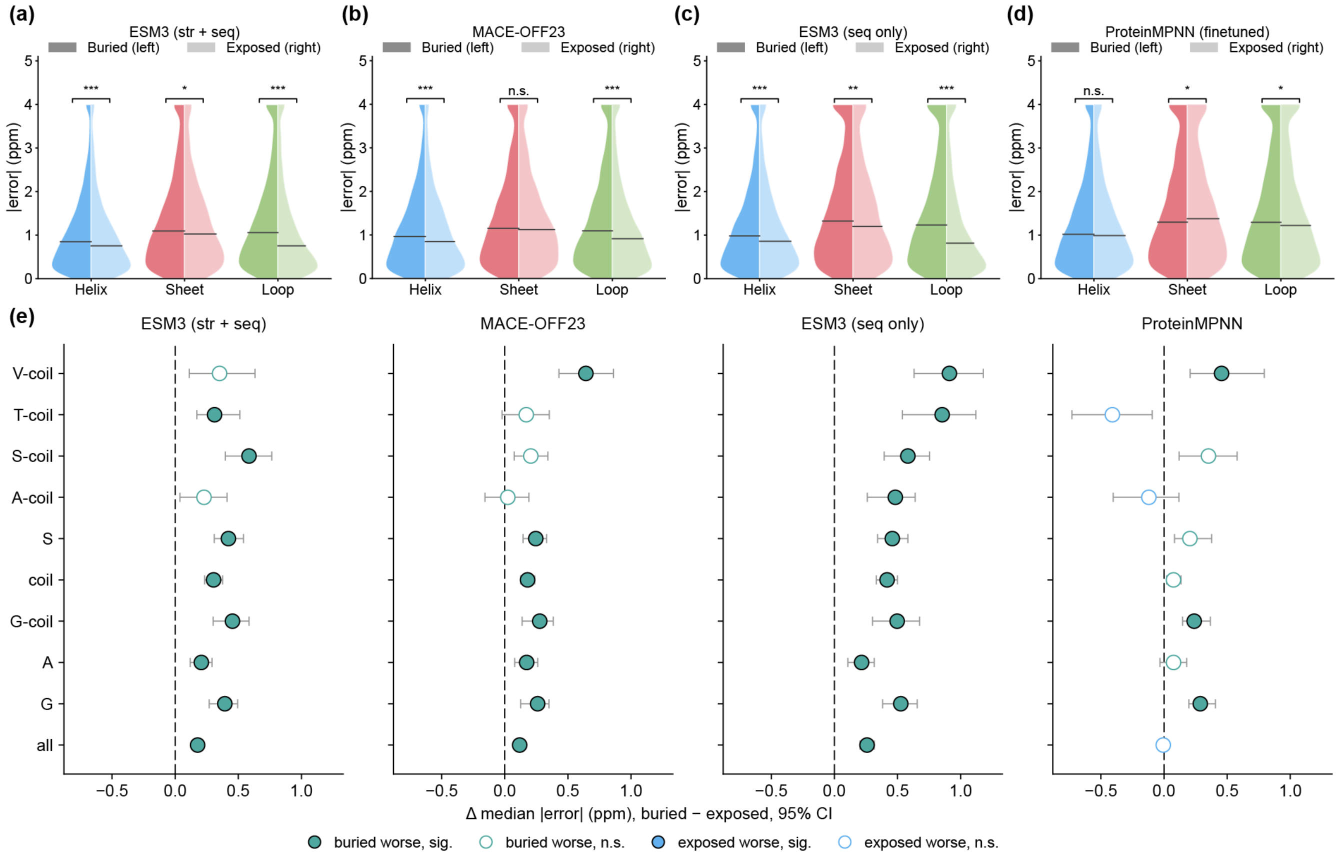


**Fig. S8 | Benchmarking SLAE-CS against predictors built on other foundation model representations.**

(a–d) $^{15}$N |error| of models by secondary structure: ESM3 (structure + sequence; a), MACE-OFF23 (b), ESM3 (sequence only; c) and fine-tuned ProteinMPNN (d). Each violin is split into buried residues (left half) and exposed residues (right half). Horizontal lines mark medians. Brackets compare buried with exposed residues (*, $p < 0.05$; **, $p < 0.01$; ***, $p < 0.001$; n.s., not significant). e, Difference in median $^{15}$N absolute prediction error between buried and exposed residues (buried − exposed) for each additional predictor, evaluated in the strata that showed a significant effect for SLAE-CS (Fig. 5e). Positive values indicate larger errors for buried residues. Teal markers denote strata in which buried residues have larger errors; blue markers denote strata in which exposed residues have larger errors. Filled markers indicate significance (Bonferroni-corrected $P < 0.01$); open markers indicate non-significance. Error bars denote 95% confidence intervals from 100,000 bootstrap resamples of protein clusters. P values were Bonferroni-corrected for the number of strata tested.

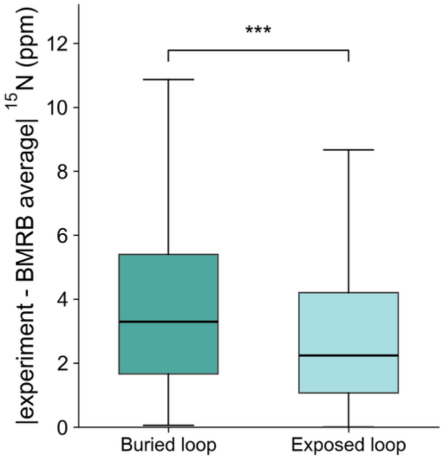


**Fig. S9 | Experimental backbone $^{15}N$ chemical shifts of buried loop residues deviate more from the residue-type database average than those of exposed loop residues.**

Box plots show the per-residue absolute deviation of the experimental backbone amide $^{15}N$ chemical shift from the BMRB average $^{15}N$ shift of the same residue type (Table S1). Residues are loop residues from the 48 test-set proteins with 1-μs MD trajectories. Boxes show the median and interquartile range. Whiskers extend to 1.5 × IQR, and outliers are not shown. The medians are 3.30 ppm (buried) and 2.24 ppm (exposed). The CI comes from a protein-level cluster bootstrap (B = 2000). ***, two-sided bootstrap $p < 5 \times 10^{-4}$.

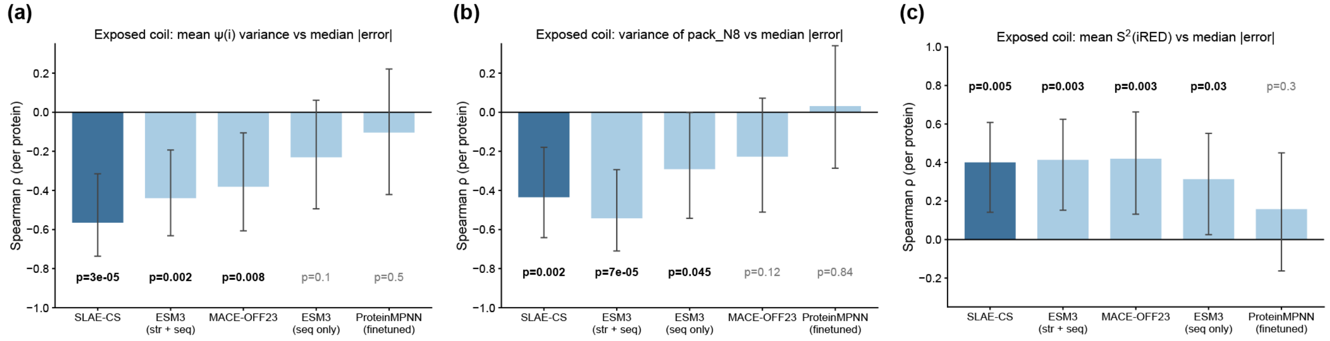


**Fig. S10 | Prediction error in exposed loops correlates with backbone dynamics and local packing across proteins.**

Spearman correlation (ρ) between per-protein median $^{15}N$ absolute prediction error for exposed loop residues and three per-protein dynamic features, shown for each predictor (n = 48 proteins). The features are: (a) circular variance of ψ(i) over MD frames; (b) variance over MD frames of pack_N8, the number of heavy atoms within 8 Å of the backbone amide nitrogen; (c) backbone N–H order parameter $S^2$ estimated by iRED. Each feature was averaged over exposed loop residues within each protein. Error bars, 95% confidence intervals from 2,000 bootstrap resampling over proteins. Values with $P < 0.05$ are shown in bold.

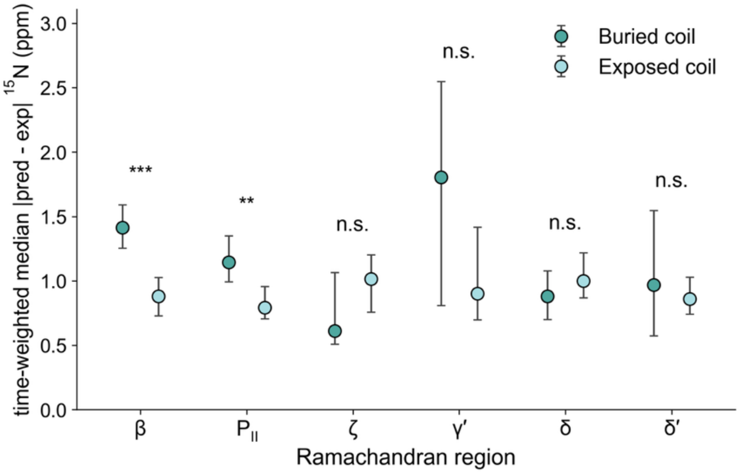


**Fig. S11 | The higher $^{15}N$ error of buried loops is confined to the β and $P_{II}$ regions of the Ramachandran map.**

Time-weighted median absolute error of SLAE-CS backbone amide $^{15}N$ predictions for buried (dark teal) and exposed (light teal) loop residues, resolved by Ramachandran region. Residues are from the 48 test-set proteins with 1-μs MD trajectories. Each frame was assigned to a Ramachandran region of Hollingsworth and Karplus (2010). Points, weighted medians; whiskers, 95% confidence intervals from a protein-level bootstrap (2,000 resamples). Asterisks, two-sided bootstrap P for the buried − exposed difference (***$P < 0.001$; **$P < 0.01$; n.s., not significant). Regions are shown only when both groups have at least 5 residue-equivalents of summed time; α and PII′ fall below this threshold for buried loops and are omitted. Buried loops are predicted worse than exposed loops in the β (1.41 vs 0.88 ppm) and PII (1.14 vs 0.79 ppm) regions, but not in the ζ, γ′, δ or δ′ regions.

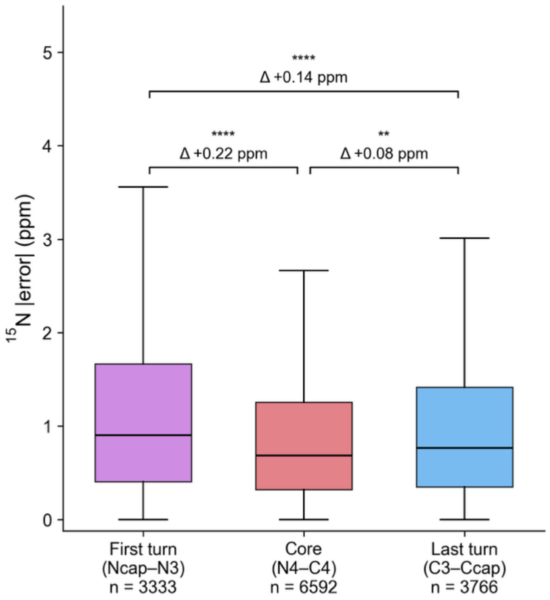


**Fig. S12 | SLAE-CS $^{15}N$ prediction errors are elevated at α-helix termini, most strongly in the first turn.**

Absolute $^{15}N$ prediction error of SLAE-CS for α-helix residues, grouped by position along the helix: first turn (Ncap–N3; n = 3,333 residues), core (N4–C4; n = 6,592) and last turn (C3–Ccap; n = 3,766). Proline residues were excluded. Boxes show the median and interquartile range (IQR); whiskers extend to 1.5 × IQR; outliers are not shown. Brackets give the difference between pooled group medians (first turn − core, last turn − core and first turn − last turn). P values are from two-sided Wilcoxon signed-rank tests on per-protein differences in median error, Holm-adjusted for three comparisons. **$P < 0.01$; ****$P < 10^{-4}$.

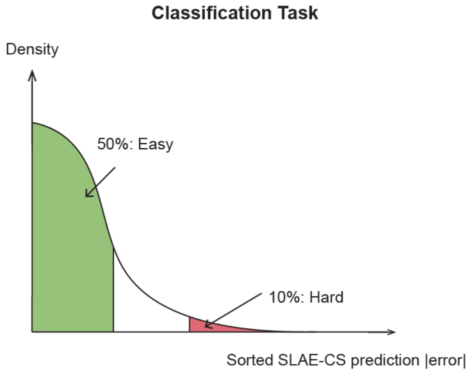


**Fig. S13 | Classification scheme for distinguishing hard-to-predict from easy-to-predict residues**

Test set predictions were ranked by |error|, with the worst 10% labeled "hard" and the best 50% labeled "easy".

### Detailed analyses of dynamics and complexity descriptors

All three descriptors here showed weak, non-monotonic association with prediction errors. Because $S^2$ reports fast picosecond-to-nanosecond motions whereas p(exchange) captures slower microsecond-to-millisecond conformational exchange, their weak inverse associations suggest that motions on different timescales contribute to distinct aspects of prediction error. However, the substantial spread within each bin indicates that considerable heterogeneity in prediction error remains unexplained by any single descriptor, motivating a model capable of integrating these and other structural features jointly rather than relying on any one univariate relationship.

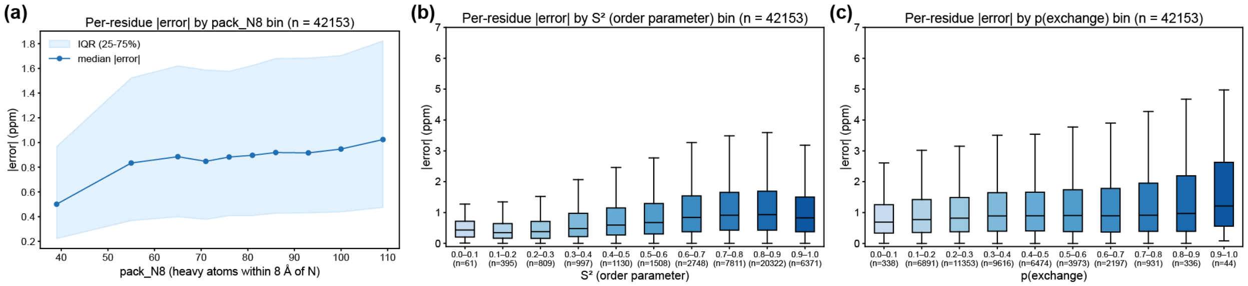


**Fig. S14 | Per-residue $^{15}$N prediction error increases modestly with local packing density, backbone order and exchange probability.**

Per-residue absolute $^{15}$N prediction error of SLAE-CS for test-set proteins. (a) Error as a function of local packing density (pack_N8, the number of heavy atoms within 8 Å of the backbone amide nitrogen. Residues were divided into ten equal-count bins; points show the median error of each bin and the shaded band shows the interquartile range (IQR). Median error rises from 0.5 ppm in the lowest bin (pack_N8 12–47) to 1.02 ppm in the highest bin (pack_N8 104–131). (b) Error binned by backbone N–H order parameter $S^2$ (equal-width bins of 0.1). Median error rises from 0.43 ppm ($S^2$ 0.0–0.1) to a maximum of 0.93 ppm ($S^2$ 0.8–0.9) and declines to 0.83 ppm in the most rigid bin ($S^2$ 0.9–1.0). (c) Error binned by p(exchange) (equal-width bins of 0.1). Median error rises from 0.69 ppm (0.0–0.1), and reaches 1.21 ppm in the sparsely populated highest bin (0.9–1.0; n = 44). Box plots in b and c: centre line, median; box, IQR; whiskers, most extreme values within 1.5 × IQR; outliers not shown. The number of residues per bin is given below each box.

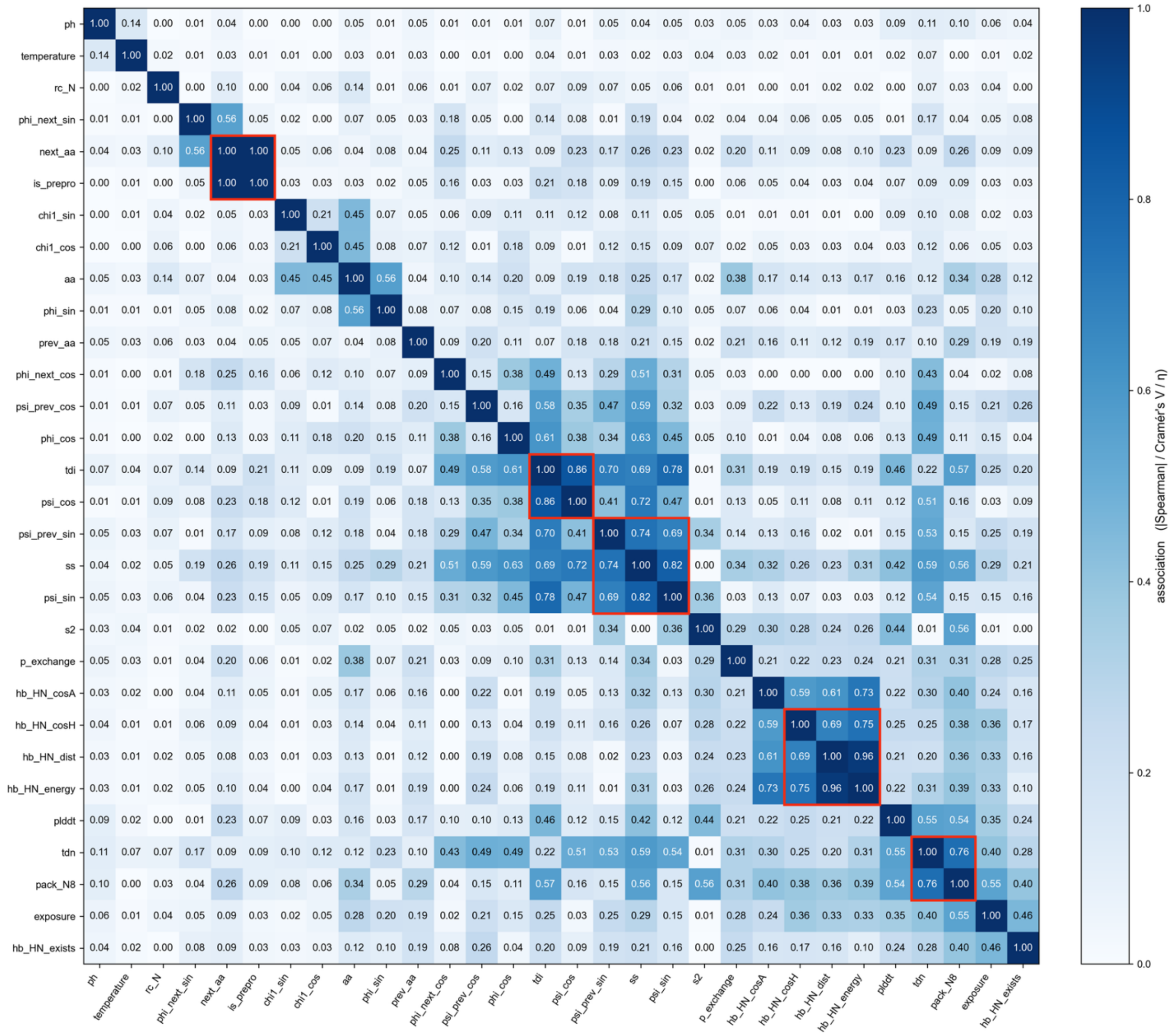


**Fig. S15 | Pairwise associations among the per-residue features used in the $^{15}$N error analysis.**

Associations between features, computed on test set. A type-aware metric is used (color bar): |Spearman ρ| for numeric–numeric pairs, bias-corrected Cramér's V for categorical–categorical pairs, and the correlation ratio η for categorical–numeric pairs. The diagonal is 1. Features are ordered by average-linkage hierarchical clustering on the distance 1 − association. Red boxes mark groups of mutually associated features obtained by cutting the dendrogram at an association threshold of 0.7; features within a box are collapsed into a single group to remove redundancy before the grouped feature-importance analysis (Fig. 6c).

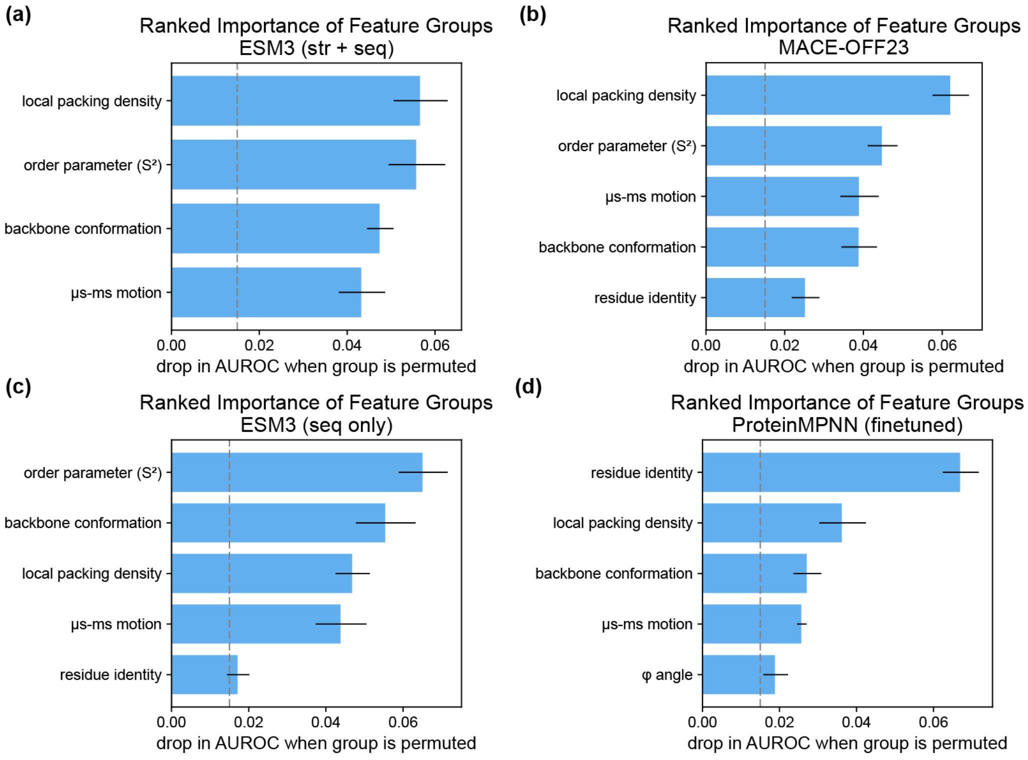


**Fig. S16 | Grouped permutation importance for the hard-versus-easy classification across other predictors.**

Same analysis as Fig. 6c, applied to the errors of each additional predictor: (a) ESM3 (structure + sequence); (b) MACE-OFF23; (c) ESM3 (sequence only); (d) fine-tuned ProteinMPNN. Importance is the decrease in AUROC when each feature group is shuffled within each of the K cross-validation folds (see Methods). Error bars, s.d. across folds. Only groups with importance > 0.015 (dashed line) are shown, ranked by importance.

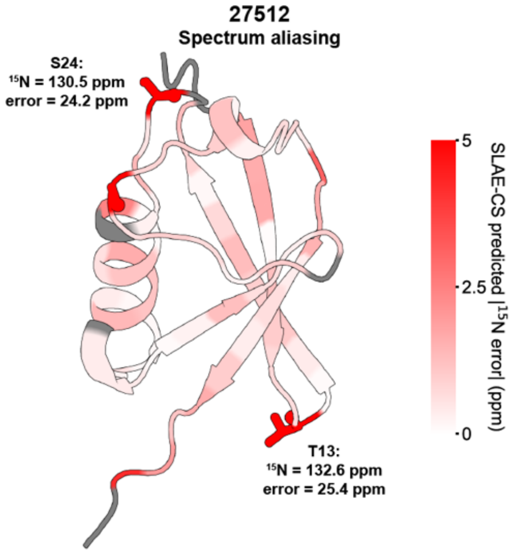


**Fig. S17 | Spectral aliasing in the deposited $^{15}N$ shifts likely accounts for the large per-protein error of SLAE-CS on ubiquitin G76C (BMRB 27512).**

AlphaFold2 structure of ubiquitin G76C colored by SLAE-CS-predicted $^{15}N$ absolute error (white, low; red, high, saturating at 5 ppm; grey, no data). Residues with absolute error > 5 ppm are shown as sticks. Two residues, T13 and S24 (shown as sticks), have deposited $^{15}N$ shifts of 132.6 and 130.5 ppm. These values are atypically downfield for Thr and Ser and are consistent with spectral aliasing. Their absolute errors (25.4 and 24.2 ppm) inflate the per-protein $^{15}N$ average error from 0.95 ppm to 1.64 ppm when these two residues are included.

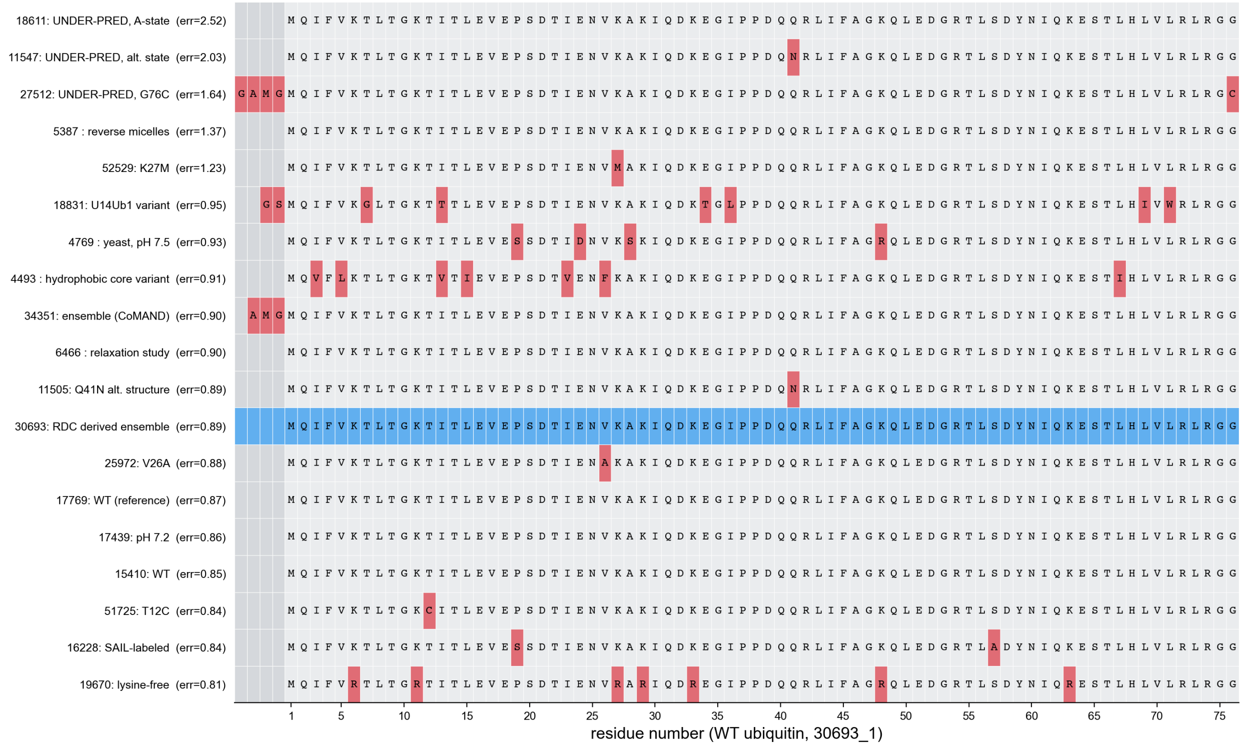


**Fig. S18 | Sequence alignment of ubiquitin entries in the test set.**

Sequences of 18 ubiquitin BMRB entries in the test set, aligned to wild-type human ubiquitin (BMRB 30693; blue). Each entry is labelled with its BMRB ID, a short description and the mean $^{15}N$ absolute prediction error of SLAE-CS in ppm (in parentheses). Entries are ordered by this error. Red cells mark positions that differ from the reference, including substitutions and N-terminal expression-tag residues; grey cells mark alignment gaps. Residue numbering on the x axis follows wild-type ubiquitin (M1–G76).

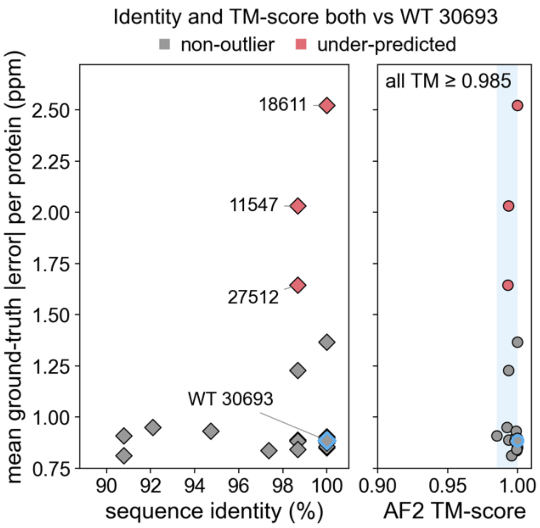


**Fig. S19 | Ubiquitin entries with near-identical inputs differ widely in SLAE-CS prediction error.**

Mean $^{15}N$ absolute prediction error of SLAE-CS for each ubiquitin entry in the test set (n = 19, including wild type), plotted against sequence identity (left) and TM-score of the AlphaFold2 input model (right). Both are computed relative to wild-type ubiquitin (BMRB 30693; blue outline). Red, under-predicted entries, labelled by BMRB ID; grey, all other entries. All entries share ≥ 90% sequence identity with wild type and have AlphaFold2 models with the wild-type fold (TM-score ≥ 0.985; shaded), yet their mean error ranges from 0.8 to 2.5 ppm.

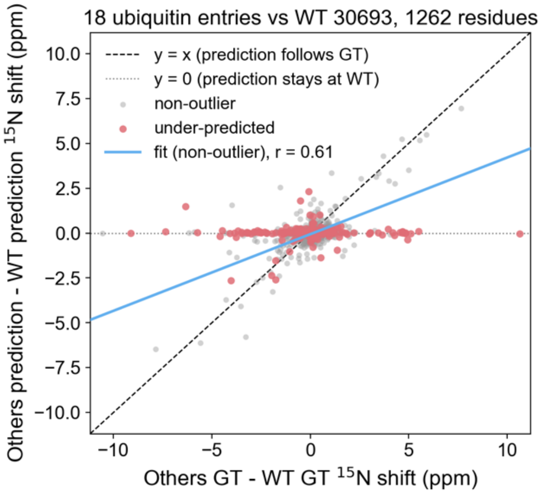


**Fig. S20 | SLAE-CS does not reproduce chemical shift differences between ubiquitin entries when the differences are not encoded in the input structure.**

Backbone $^{15}N$ shifts of 18 ubiquitin entries, expressed relative to wild-type ubiquitin (BMRB 30693). The x axis shows the experimental shift minus the wild-type experimental shift; the y axis shows the SLAE-CS-predicted shift minus the wild-type predicted shift. On the dashed line (y = x), the prediction reproduces the experimental change. On the dotted line (y = 0), the prediction is unchanged from wild type. Red, residues from the under-predicted entries (18611, 11547 and 27512). Grey, all other entries. The blue line is a least-squares fit to the grey points (Pearson r = 0.61, slope=0.43). Residues with | experimental $\Delta^{15}N$| > 15 ppm relative to wild type were excluded; these may be attributable to spectral aliasing in the deposited data.

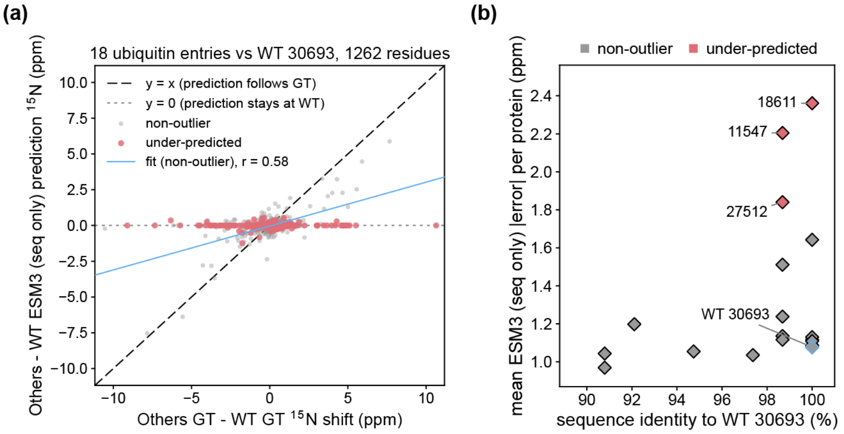


**Fig. S21 | ESM3 (sequence only) cannot reproduce chemical shift differences between ubiquitin entries.**

Here ESM3 (sequence only) stands for predictor based on it. (a) Backbone $^{15}$N shifts of 18 ubiquitin entries, expressed relative to wild-type ubiquitin (BMRB 30693). The x axis shows the experimental shift minus the wild-type experimental shift; the y axis shows the shift predicted by ESM3 (sequence only) minus the wild-type predicted shift. On the dashed line (y = x), the prediction reproduces the experimental change. On the dotted line (y = 0), the prediction is unchanged from wild type. Red, residues from the under-predicted entries (18611, 11547 and 27512). Grey, all other entries. The blue line is a least-squares fit to the grey points (Pearson r = 0.58, slope = 0.31). Residues with |experimental $\Delta^{15}$N| > 15 ppm relative to wild type were excluded, as these likely reflect spectral aliasing in the deposited data. (b) Mean $^{15}$N absolute prediction error of ESM3 (sequence only) for each ubiquitin entry, plotted against sequence identity relative to wild type (BMRB 30693; blue outline). Under-predicted entries are labelled. All entries share ≥ 90% sequence identity with wild type, yet their mean error ranges from 0.97 to 2.36 ppm.

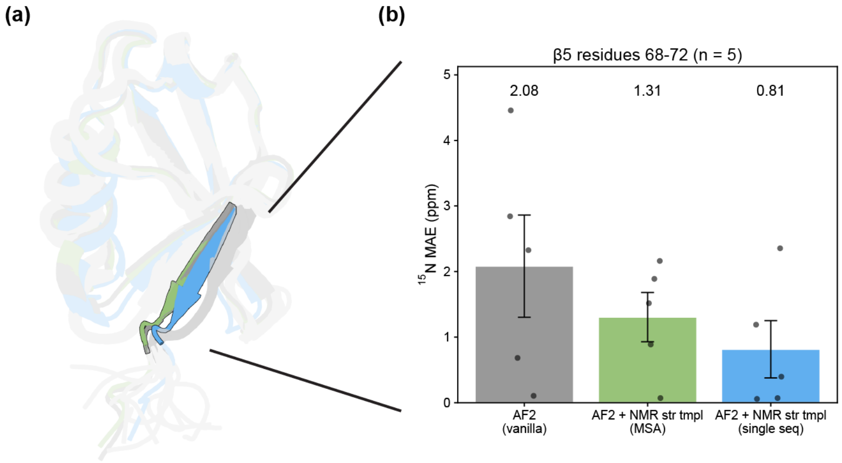


**Fig. S22 | Steering the input structure toward the NMR structure reduces SLAE-CS $^{15}$N error in β5 of the ubiquitin N2 state.**

(a) Superposition of three AF2 input structures for ubiquitin Q41N, which populates the alternative N2 state (BMRB 11547). The three structures are: the default AF2 model used in the test set (vanilla; grey); AF2 with the deposited NMR structure (PDB 2RU6, model 1) as the only template and a full MSA (green); and AF2 with the same template in single-sequence mode, which forces AF2 to rely on the template (blue). β5 (residues 68–72) is highlighted; the remainder of each structure is shown transparent. (b) SLAE-CS $^{15}$N mean absolute error over β5 residues (n = 5) for each input structure. Bars show the MAE of predictions (2.08, 1.31 and 0.81 ppm); dots show individual residues; error bars indicate the standard error of the mean across the five β5 residues.

Table S1. BMRB backbone chemical-shift statistics used for outlier screening. Mean and standard deviation (s.d.) of backbone $^{1}$H, $^{15}$N, $^{13}$C$^{\alpha}$, $^{13}$C$^{\beta}$ and $^{13}$C′ chemical shifts (p.p.m.) for the 20 standard amino acids.

| | $^{1}$H | | $^{15}$N | | $^{13}$C$^{\alpha}$ | | $^{13}$C$^{\beta}$ | | $^{13}$C′ | |
|---|---|---|---|---|---|---|---|---|---|---|
| **Residue** | **Mean** | **s.d.** | **Mean** | **s.d.** | **Mean** | **s.d.** | **Mean** | **s.d.** | **Mean** | **s.d.** |
| Ala | 8.197 | 0.878 | 123.395 | 5.602 | 53.132 | 2.603 | 19.023 | 2.828 | 177.742 | 3.443 |
| Arg | 8.240 | 0.971 | 120.871 | 4.520 | 56.756 | 3.136 | 30.664 | 2.374 | 176.423 | 3.129 |
| Asn | 8.353 | 1.864 | 118.945 | 5.194 | 53.517 | 3.230 | 38.704 | 3.282 | 175.208 | 3.308 |
| Asp | 8.295 | 0.579 | 120.727 | 4.449 | 54.664 | 2.578 | 40.878 | 2.391 | 176.354 | 3.351 |
| Cys | 8.372 | 0.693 | 120.340 | 16.415 | 57.969 | 3.489 | 33.451 | 6.662 | 174.723 | 3.256 |
| Gln | 8.243 | 1.806 | 120.039 | 4.327 | 56.518 | 2.507 | 29.171 | 2.372 | 176.306 | 8.487 |
| Glu | 8.331 | 0.786 | 120.763 | 4.432 | 57.303 | 3.061 | 29.987 | 2.908 | 176.830 | 3.952 |
| Gly | 8.327 | 0.851 | 109.636 | 6.387 | 45.360 | 2.055 | – | – | 173.831 | 3.460 |
| His | 8.258 | 1.202 | 119.695 | 5.022 | 56.482 | 3.212 | 30.318 | 2.994 | 175.128 | 4.436 |
| Ile | 8.256 | 0.686 | 121.431 | 5.824 | 61.627 | 3.231 | 38.553 | 2.764 | 175.832 | 4.220 |
| Leu | 8.218 | 0.718 | 121.962 | 7.212 | 55.643 | 2.202 | 42.209 | 1.987 | 177.010 | 3.415 |
| Lys | 8.190 | 1.483 | 121.069 | 4.699 | 56.935 | 3.015 | 32.762 | 2.728 | 176.621 | 5.198 |
| Met | 8.258 | 1.147 | 120.152 | 4.893 | 56.112 | 2.265 | 32.954 | 3.016 | 176.192 | 3.140 |
| Phe | 8.332 | 0.727 | 120.401 | 5.403 | 58.093 | 3.591 | 39.901 | 3.346 | 175.441 | 2.863 |
| Pro | – | – | – | – | 63.319 | 3.304 | 31.871 | 2.868 | 176.669 | 3.918 |
| Ser | 8.277 | 0.685 | 116.321 | 4.059 | 58.664 | 3.336 | 63.721 | 4.455 | 174.547 | 3.511 |
| Thr | 8.226 | 0.630 | 115.383 | 6.022 | 62.193 | 2.705 | 69.607 | 5.071 | 174.453 | 3.796 |
| Trp | 8.264 | 0.783 | 121.634 | 5.828 | 57.727 | 4.422 | 30.030 | 4.376 | 176.040 | 5.442 |
| Tyr | 8.282 | 0.887 | 120.648 | 10.623 | 58.126 | 2.967 | 39.267 | 2.948 | 175.406 | 4.256 |
| Val | 8.267 | 0.672 | 121.154 | 6.848 | 62.483 | 3.122 | 32.689 | 2.189 | 175.639 | 3.203 |

Table S2. Entry pairs kept for the intrinsic noise analyses

| No. | Entry A | Entry B | PANAV intrinsic RMSE (ppm) | LACS intrinsic RMSE (ppm) |
|---|---|---|---|---|
| 1 | bmr19458_1 | bmr6149_1 | 7.884 | 8.112 |
| 2 | bmr17836_1 | bmr17837_1 | 4.189 | 4.628 |
| 3 | bmr18458_1 | bmr30098_1 | 4.161 | 4.156 |
| 4 | bmr27185_1 | bmr52587_1 | 3.645 | 3.720 |
| 5 | bmr17085_1 | bmr18458_1 | 3.673 | 3.670 |
| 6 | bmr4765_1 | bmr11451_1 | 3.298 | 3.461 |
| 7 | bmr6321_1 | bmr6322_1 | 1.982 | 2.038 |
| 8 | bmr17590_1 | bmr17591_1 | 1.913 | 1.901 |
| 9 | bmr31112_1 | bmr31113_1 | 1.776 | 1.769 |
| 10 | bmr30432_1 | bmr30434_1 | – | 1.715 |
| 11 | bmr31107_1 | bmr52019_1 | 1.587 | 1.570 |
| 12 | bmr19461_1 | bmr19462_1 | 1.544 | 1.432 |
| 13 | bmr50621_1 | bmr50622_1 | 1.486 | 1.514 |
| 14 | bmr30172_1 | bmr34039_1 | 1.468 | 1.492 |
| 15 | bmr6356_1 | bmr19991_1 | 1.365 | 1.383 |
| 16 | bmr4031_1 | bmr19065_1 | 1.293 | – |
| 17 | bmr25225_1 | bmr25226_1 | 0.867 | 1.233 |
| 18 | bmr25226_1 | bmr34547_1 | 0.866 | 1.233 |
| 19 | bmr25225_1 | bmr34548_1 | 0.862 | 1.233 |
| 20 | bmr34547_1 | bmr34548_1 | 0.861 | 1.233 |
| 21 | bmr15558_1 | bmr17477_1 | 0.890 | 1.139 |
| 22 | bmr2060_1 | bmr2371_1 | 1.134 | – |
| 23 | bmr17085_1 | bmr30098_1 | 1.124 | 1.122 |
| 24 | bmr17357_1 | bmr25350_1 | 0.119 | 1.108 |

| No. | Entry A | Entry B | PANAV intrinsic RMSE (ppm) | LACS intrinsic RMSE (ppm) |
|---|---|---|---|---|
| 25 | bmr26635_1 | bmr30825_1 | 1.086 | – |
| 26 | bmr4023_1 | bmr5656_1 | 0.350 | 1.065 |
| 27 | bmr30331_1 | bmr30332_1 | 0.634 | 1.034 |
| 28 | bmr50010_1 | bmr50011_1 | 0.628 | 0.930 |
| 29 | bmr6911_1 | bmr16558_1 | 0.327 | 0.903 |
| 30 | bmr6317_1 | bmr17892_1 | 0.826 | 0.813 |
| 31 | bmr16142_1 | bmr16143_1 | 0.219 | 0.790 |
| 32 | bmr27663_1 | bmr30531_1 | 0.097 | 0.710 |
| 33 | bmr27497_1 | bmr30471_1 | 0.658 | 0.695 |
| 34 | bmr27239_2 | bmr27242_1 | 0.676 | – |
| 35 | bmr30419_1 | bmr30420_1 | 0.636 | – |
| 36 | bmr6732_1 | bmr6734_1 | 0.629 | – |
| 37 | bmr15894_1 | bmr15895_1 | 0.566 | 0.568 |
| 38 | bmr15330_1 | bmr15358_1 | 0.534 | 0.545 |
| 39 | bmr15332_1 | bmr15355_1 | 0.526 | 0.524 |
| 40 | bmr30394_1 | bmr30395_1 | 0.523 | 0.522 |
| 41 | bmr6732_1 | bmr6735_1 | 0.505 | – |
| 42 | bmr19240_1 | bmr19241_1 | 0.483 | 0.488 |
| 43 | bmr19323_1 | bmr19324_1 | 0.456 | 0.459 |
| 44 | bmr15149_1 | bmr15502_1 | 0.420 | – |
| 45 | bmr6734_1 | bmr6735_1 | 0.395 | – |
| 46 | bmr17074_1 | bmr17076_1 | 0.349 | 0.348 |
| 47 | bmr4098_1 | bmr15429_1 | 0.297 | 0.279 |
| 48 | bmr15689_1 | bmr16895_1 | 0.259 | 0.026 |
| 49 | bmr25272_1 | bmr34383_1 | 0.182 | 0.122 |
| 50 | bmr16818_1 | bmr16848_1 | 0.176 | 0.149 |
| 51 | bmr51744_1 | bmr51746_1 | 0.172 | – |
| 52 | bmr17109_1 | bmr18152_1 | 0.141 | 0.164 |
| 53 | bmr18327_1 | bmr18832_1 | 0.152 | 0.142 |
| 54 | bmr34803_1 | bmr34918_1 | 0.148 | 0.026 |
| 55 | bmr51745_1 | bmr51746_1 | 0.132 | – |
| 56 | bmr34160_1 | bmr34164_1 | 0.119 | 0.119 |
| 57 | bmr19406_1 | bmr19412_1 | 0.118 | – |
| 58 | bmr6249_1 | bmr15158_1 | 0.112 | – |
| 59 | bmr6140_1 | bmr15607_1 | 0.103 | 0.082 |
| 60 | bmr16215_1 | bmr5723_1 | 0.099 | – |
| 61 | bmr16805_1 | bmr18263_1 | 0.084 | 0.078 |
| 62 | bmr25226_1 | bmr34548_1 | 0.082 | – |
| 63 | bmr51769_1 | bmr51770_1 | 0.074 | – |
| 64 | bmr18777_1 | bmr19952_1 | 0.065 | 0.066 |
| 65 | bmr5917_1 | bmr10023_1 | 0.061 | – |
| 66 | bmr51744_1 | bmr51745_1 | 0.060 | – |
| 67 | bmr50838_1 | bmr50963_1 | 0.059 | – |
| 68 | bmr5270_1 | bmr5758_1 | 0.053 | 0.046 |
| 69 | bmr16908_1 | bmr17805_1 | 0.043 | – |
| 70 | bmr17820_1 | bmr25113_1 | 0.034 | 0.029 |
| 71 | bmr16272_1 | bmr16283_1 | 0.032 | 0.032 |
| 72 | bmr25225_1 | bmr34547_1 | 0.012 | – |

Table S3. Entry pairs manually excluded for the intrinsic noise analyses

| No. | Entry A | Entry B | Category | Reason for exclusion | PANAV intrinsic RMSE (ppm) | LACS intrinsic RMSE (ppm) |
|---|---|---|---|---|---|---|
| 1 | bmr11035_1 | bmr11037_1 | Cellular environment | in vitro vs in-cell (M9 slurry, confirmed) | 0.155 | 0.167 |
| 2 | bmr11049_1 | bmr11050_1 | Membrane-mimetic (micelle) | DPC/SDS micelle | 1.105 | – |
| 3 | bmr11505_1 | bmr11547_1 | Point mutation / pressure | Q41N mutant at 1 atm vs wild type at 2500 bar | 6.214 | 6.290 |
| 4 | bmr11519_1 | bmr11520_1 | Molecular crowding | dilute vs crowded condition | 0.398 | – |
| 5 | bmr11519_1 | bmr11521_1 | Molecular crowding | crowding (protein-G B1 fusion crowding agent) | 0.377 | – |
| 6 | bmr11520_1 | bmr11521_1 | Molecular crowding | Villin crowded vs crowded+protein G (different crowding agent) | 0.046 | – |
| 7 | bmr12035_1 | bmr12038_1 | Phosphorylation | phosphorylated | 0.582 | 0.630 |
| 8 | bmr12036_1 | bmr12039_1 | Phosphorylation | phosphorylated | 0.315 | 0.310 |
| 9 | bmr15176_1 | bmr15865_1 | Point mutation | different point mutations (D5G,C72S vs C72S) | 0.320 | 0.286 |
| 10 | bmr15264_1 | bmr16265_1 | Redox state | redox | 0.434 | 0.434 |
| 11 | bmr15280_1 | bmr6709_1 | Redox state | oxidized (Cys67–Cys70 disulfide) vs reduced form | 1.591 | 1.586 |
| 12 | bmr15599_1 | bmr6625_1 | Ligand/partner binding | Pi free and bound form | 5.712 | 5.633 |
| 13 | bmr15930_1 | bmr15936_1 | Cosolvent (TFE) | sigma70 region4, 10% TFE vs 0-30% TFE series (same group as the entry below) | 1.134 | – |
| 14 | bmr15930_1 | bmr15975_1 | Cosolvent (TFE) | 10% vs 30% TFE | 0.933 | 0.928 |
| 15 | bmr15936_1 | bmr15975_1 | Cosolvent (TFE) | sigma70 region4, 0-30% TFE gradient series vs 30% TFE | 1.471 | – |
| 16 | bmr16166_1 | bmr16800_1 | Cosolvent (TFE) | TFE difference | 1.183 | 1.827 |
| 17 | bmr16275_1 | bmr20082_1 | Membrane-mimetic (micelle) | micelle: DPC vs SDS | – | 0.865 |
| 18 | bmr16445_1 | bmr26554_1 | Membrane-mimetic (micelle) | SDS micelle | 2.308 | 2.278 |
| 19 | bmr16603_1 | bmr7432_1 | Ligand/partner binding | apo-IscU(D39A) vs IscU(D39A) "interaction with HscB" | 0.055 | 0.011 |
| 20 | bmr16798_1 | bmr16799_1 | Cosolvent (TFE) | TFE difference | 1.015 | 1.621 |
| 21 | bmr17701_1 | bmr18037_1 | Covalent/PTM modification | L,D-transpeptidase unmodified vs imipenem-acylated (antibiotic covalent adduct) | 0.474 | 0.473 |
| 22 | bmr17724_1 | bmr18740_1 | Membrane-mimetic (micelle) | SDS vs LMPG micelles | 0.920 | 1.029 |
| 23 | bmr1812_1 | bmr1813_1 | Redox state | redox state (reduced/oxidized thioredoxin) | 0.470 | – |
| 24 | bmr18224_1 | bmr18226_1 | Membrane-mimetic (micelle) | SDS/LPPG micelle | 0.692 | 0.700 |
| 25 | bmr18412_1 | bmr30614_1 | Ligand/partner binding | free PEA-15 vs PEA-15 in a 1:1 complex with unlabeled ERK2 (ERK2 not listed in the BMRB sample; Int. J. Mol. Sci. 2019, 20, 3335) | 0.751 | 1.047 |
| 26 | bmr18567_1 | bmr18969_1 | Point mutation | different point-mutation sites (S114A vs S55A) | 0.373 | 0.723 |
| 27 | bmr18630_1 | bmr18631_1 | Redox state | redox | 0.337 | 0.366 |
| 28 | bmr18679_1 | bmr18680_1 | Ligand/partner binding | dT7 difference | 0.804 | 0.800 |
| 29 | bmr19189_1 | bmr25333_1 | Lipid environment | different lipids | 0.918 | 1.240 |
| 30 | bmr19480_1 | bmr19481_1 | Phosphorylation | phosphorylation | 0.117 | 0.101 |
| 31 | bmr19615_1 | bmr19616_1 | Ligand/partner binding | apo vs L-citrulline-bound R40E/R98H/C249S PaDDAH (Rasheed et al. Biochemistry 2014) | 0.812 | 0.813 |

| No. | Entry A | Entry B | Category | Reason for exclusion | PANAV intrinsic RMSE (ppm) | LACS intrinsic RMSE (ppm) |
|---|---|---|---|---|---|---|
| 32 | bmr19742_1 | bmr19743_1 | Ligand/partner binding | L-citrulline-bound vs apo R40E/R98H PaDDAH (Rasheed et al. Biochemistry 2014) | 2.720 | 2.693 |
| 33 | bmr19947_1 | bmr19963_1 | Disulfide isomer | different disulfide connectivity (C5–C24, C16–C34, C18–C29 vs C5–C34, C16–C29, C18–C24) | 3.872 | 3.895 |
| 34 | bmr25179_1 | bmr25586_1 | Membrane-mimetic (micelle) | SDS micelles vs aqueous solution | 1.516 | – |
| 35 | bmr25180_1 | bmr25676_1 | Membrane-mimetic (micelle) | SDS micelles vs aqueous solution | 1.685 | 1.654 |
| 36 | bmr25602_1 | bmr26574_1 | Ligand/partner binding | binding partner | 0.968 | 0.978 |
| 37 | bmr25632_1 | bmr26573_1 | Ligand/partner binding | binding partner GAT | 3.058 | 2.585 |
| 38 | bmr25663_1 | bmr6392_1 | Ligand/partner binding | binding | 0.571 | 0.570 |
| 39 | bmr25817_1 | bmr25818_1 | Membrane-mimetic (micelle) | SDS/DPC micelle | 0.631 | 0.629 |
| 40 | bmr25875_1 | bmr25879_1 | Ligand/partner binding | binding | 0.780 | 1.336 |
| 41 | bmr25909_1 | bmr25910_1 | Cellular environment | in vitro vs in-cell | 0.232 | 0.098 |
| 42 | bmr26672_1 | bmr27887_1 | Phase state (LLPS) | FUS LC dispersed phase condensed phase | 0.233 | – |
| 43 | bmr26746_1 | bmr30077_1 | Ligand/partner binding | GMPPNP-bound vs apo | 0.779 | – |
| 44 | bmr26893_1 | bmr26894_1 | Molecular crowding | crowding; component list confirms lysozyme was added | 0.398 | 0.854 |
| 45 | bmr26967_1 | bmr26968_1 | Glycosylation | glycosylation (GlcNAc) difference | 0.264 | 0.264 |
| 46 | bmr27237_1 | bmr27238_1 | Covalent/PTM modification | S-nitrosylated | 0.338 | 0.343 |
| 47 | bmr27269_1 | bmr30312_1 | Redox state | TCEP-reduced (10 mM TCEP, disulfides broken) vs oxidized disulfide-bonded gHEEE_02 | 3.063 | 3.050 |
| 48 | bmr27365_1 | bmr30435_1 | Membrane-mimetic (micelle) | 250 mM DPC micelle vs 20% TFE | – | 0.804 |
| 49 | bmr27396_1 | bmr30404_1 | Ligand/partner binding | myo-inositol hexakisphosphate binding | 0.487 | 0.491 |
| 50 | bmr27543_1 | bmr27709_1 | Membrane-mimetic (micelle) | different micelles (LMPG vs TDPC:SDS) | 0.633 | 0.330 |
| 51 | bmr27721_2 | bmr27722_2 | Ligand stereoisomer | bound to a different ligand (compound S vs R) | 0.665 | – |
| 52 | bmr27915_1 | bmr27916_1 | Oligomerization state | homodimer vs heterodimer (component list has an extra p65) | 0.384 | 0.308 |
| 53 | bmr28008_1 | bmr28009_1 | Ligand/partner binding | binding GDP vs GMPPNP | 1.153 | 1.394 |
| 54 | bmr30234_1 | bmr52078_1 | Ubiquitin linkage isomer | K27- vs K6-linked diubiquitin | 0.625 | – |
| 55 | bmr30234_1 | bmr52108_2 | Ubiquitin linkage isomer | K27- vs K29-linked diubiquitin | 0.970 | – |
| 56 | bmr30234_1 | bmr52115_2 | Ubiquitin linkage isomer | K27- vs K33-linked diubiquitin | 0.969 | – |
| 57 | bmr30234_1 | bmr52139_1 | Ubiquitin linkage isomer | K27- vs K48-linked diubiquitin | 0.595 | – |
| 58 | bmr30416_1 | bmr30435_1 | Membrane-mimetic (micelle) | DPC micelles vs 20% TFE | – | 0.804 |
| 59 | bmr30662_1 | bmr30664_1 | Point mutation | WT vs I24N point mutant | 2.251 | 2.091 |

| No. | Entry A | Entry B | Category | Reason for exclusion | PANAV intrinsic RMSE (ppm) | LACS intrinsic RMSE (ppm) |
|---|---|---|---|---|---|---|
| 60 | bmr30979_1 | bmr31111_1 | Disulfide isomer | NCR13, different disulfide-linkage isomers (C4:C10,C15:C30,C23:C28 vs C4:C23,C15:C30,C10:C28) | 2.954 | 2.987 |
| 61 | bmr34344_1 | bmr50362_1 | Redox state | redox | 1.218 | 1.440 |
| 62 | bmr36109_1 | bmr36110_1 | Ligand/partner binding | PSTD peptide + polysialic acid (PolySia) present vs absent (ligand binding) | – | 0.132 |
| 63 | bmr4197_1 | bmr4209_1 | Membrane-mimetic (micelle) | different micelles | 0.631 | 0.843 |
| 64 | bmr50198_1 | bmr50199_1 | Cellular environment | VAMP2 in SH-SY5Y cells vs HEK-293T cells (different cell-line intracellular environment) | 0.085 | – |
| 65 | bmr50484_1 | bmr50485_1 | Redox state | Redox | 0.235 | 0.592 |
| 66 | bmr50582_1 | bmr50583_1 | Ligand/partner binding | SIB1 vs. SIB1 + WRKY33 | 0.659 | 0.452 |
| 67 | bmr50676_1 | bmr50679_1 | Covalent/PTM modification | PPACK-thrombin vs S195M (PPACK is a covalent, irreversible active-site inhibitor); binding | 0.510 | 0.486 |
| 68 | bmr50835_1 | bmr51710_1 | Phosphorylation | phosphorylation | 0.545 | 0.404 |
| 69 | bmr50861_1 | bmr50862_1 | Redox state | redox | 1.292 | 1.416 |
| 70 | bmr50863_1 | bmr50864_1 | Redox state | oxidized vs TCEP-reduced (10 mM TCEP) Tim12 | 1.557 | 1.648 |
| 71 | bmr50895_1 | bmr50896_1 | Covalent/PTM modification | modification | 0.409 | 0.433 |
| 72 | bmr51018_1 | bmr51019_1 | Phosphorylation | phosphorylated | 0.373 | 0.308 |
| 73 | bmr51063_1 | bmr51064_1 | Ion binding | NaKd18 in 100 mM K+ vs 600 mM Na+ (ion-dependent structure) | 0.354 | – |
| 74 | bmr51269_1 | bmr51280_1 | Lipidation (myristoylation) | myristoylated vs non-myristoylated | 1.764 | – |
| 75 | bmr51270_1 | bmr51271_1 | Lipidation (myristoylation) | myristoylated vs non-myristoylated | 1.856 | – |
| 76 | bmr51272_1 | bmr51273_1 | Lipidation (myristoylation) | myristoylated vs non-myristoylated | 0.596 | – |
| 77 | bmr51274_1 | bmr51275_1 | Lipidation (myristoylation) | myristoylated vs non-myristoylated | 0.958 | – |
| 78 | bmr51276_1 | bmr51277_1 | Lipidation (myristoylation) | non-myristoylated vs myristoylated | 0.978 | – |
| 79 | bmr51278_1 | bmr51279_1 | Lipidation (myristoylation) | non-myristoylated vs myristoylated | 0.493 | – |
| 80 | bmr51305_1 | bmr51306_1 | Redox state | redox: reduced vs "neutral" (component list confirms DTT present) | 0.160 | – |
| 81 | bmr51306_1 | bmr51307_1 | Redox state | redox: "neutral" vs oxidized (component list confirms H2O2 present) | 0.047 | – |
| 82 | bmr51314_1 | bmr51315_1 | Methylation | methylated | 0.167 | 0.140 |
| 83 | bmr51919_1 | bmr51938_1 | Ligand/partner binding | IL-1β apo vs in the presence of the small-molecule antagonist S-2 | 1.325 | 1.282 |
| 84 | bmr52078_1 | bmr52108_2 | Ubiquitin linkage isomer | K6- vs K29-linked diubiquitin | 0.749 | – |
| 85 | bmr52078_1 | bmr52115_2 | Ubiquitin linkage isomer | K6- vs K33-linked diubiquitin | 0.748 | – |
| 86 | bmr52078_1 | bmr52139_1 | Ubiquitin linkage isomer | K6- vs K48-linked diubiquitin | 0.215 | – |
| 87 | bmr52108_2 | bmr52115_2 | Ubiquitin linkage isomer | K29- vs K33-linked diubiquitin | 0.014 | – |

| No. | Entry A | Entry B | Category | Reason for exclusion | PANAV intrinsic RMSE (ppm) | LACS intrinsic RMSE (ppm) |
|---|---|---|---|---|---|---|
| 88 | bmr52108_2 | bmr52139_1 | Ubiquitin linkage isomer | K29- vs K48-linked diubiquitin | 0.752 | – |
| 89 | bmr52115_2 | bmr52139_1 | Ubiquitin linkage isomer | K33- vs K48-linked diubiquitin | 0.751 | – |
| 90 | bmr52351_1 | bmr52352_1 | Substrate variant (acyl chain length) | different acyl chain-length substrate (C6 vs C8) | 0.344 | – |
| 91 | bmr52351_1 | bmr52353_1 | Substrate variant (acyl chain length) | different acyl chain-length substrate (C6 vs C10) | 0.458 | – |
| 92 | bmr52352_1 | bmr52353_1 | Substrate variant (acyl chain length) | different acyl chain-length substrate (C8 vs C10) | 0.152 | – |
| 93 | bmr52545_1 | bmr52546_1 | Membrane-mimetic (micelle) | AB37* with 20% dPC present vs free in solution | 3.320 | 3.295 |
| 94 | bmr5465_1 | bmr6158_1 | Membrane-mimetic (micelle) | detergent micelle | 1.893 | 1.907 |
| 95 | bmr6603_1 | bmr6847_1 | Redox state | redox | 1.259 | 1.255 |
